# The CspC:CspA heterodimer transduces germinant and co-germinant signals during *Clostridioides difficile* spore germination

**DOI:** 10.1101/2025.08.01.668244

**Authors:** Morgan E. McNellis, Gonzalo González-Del Pino, Juan A. Serrano-Jiménez, Emily R. Forster, A. Ioana Stoica, Ekaterina E. Heldwein, Aimee Shen

## Abstract

The clinically significant pathogen *Clostridioides difficile,* which lacks the transmembrane nutrient germinant receptors conserved in almost all spore-forming bacteria, initiates spore germination by a unique mechanism that requires two signals: a bile acid germinant and an amino acid or divalent cation co-germinant signal. While two soluble pseudoproteases, CspC and CspA, were initially identified as the germinant and co-germinant receptors, respectively, in *C. difficile*, we previously identified residues in an unstructured region of CspC that regulate the sensitivity of *C. difficile* spores to both signals. However, the mechanism by which CspC transduces these signals remained unclear. Here, we demonstrate that CspC forms a stable complex with CspA and determine the crystal structure of the CspC:CspA heterodimer. The structure reveals extensive interactions along the binding interface, including direct interactions between the unstructured region of CspC and CspA. Using structure-function analyses, we identify CspC:CspA interactions that regulate the sensitivity of *C. difficile* spores to germinant signals and show for the first time that CspA regulates *C. difficile* response to not only co-germinant but also germinant signals. While we also show that CspA can form a homodimer and determine its crystal structure, CspA homodimerization appears unimportant for *C. difficile* spore germination. Collectively, our analyses establish the CspC:CspA heterodimer, rather than its individual constituents, as a critical signaling node for sensing both germinant and co-germinant signals. They also suggest a new mechanistic model for how *C. difficile* transduces germinant signals, which could guide the development of therapeutics against this important pathogen.

## Introduction

*Clostridioides difficile* is a Gram-positive, spore-forming gastrointestinal pathogen that is a leading cause of hospital-acquired infections in many countries around the world^1–4^. Since *C. difficile* is an obligate anaerobe, its aerotolerant spores are its major transmissive form^5–7^. Accordingly, *C. difficile* infections begin when spores are ingested and encounter germinants in the gut that signal an environment favorable for *C. difficile* growth^5,8–10^. Germinant sensing initiates spore germination, whereby metabolically dormant spores degrade their protective cortex layer and transform into metabolically active vegetative cells^5,8^. Germination is thus essential for the ability of *C. difficile* to colonize hosts and establish infection.

*C. difficile* spore germination differs markedly from most spore-forming bacteria because *C. difficile* does not require nutrient signals, such as amino acids or sugars, to initiate germination^5,8^. Instead, *C. difficile* germinates in response to gut-specific bile acids, particularly taurocholate (TA)^5,8,9^. In addition to this germinant signal, *C. difficile* spores require a second, co-germinant signal in the form of an amino acid or divalent cation, typically glycine or calcium^9,11–13^. Furthermore, *C. difficile* is only one of two spore-forming organisms that do not encode the transmembrane Ger receptors used to sense amino acid and sugar germinant signals^14^. Ger receptors are nutrient-gated channels^15^ that bind to their nutrient germinant signals and initiate a signaling cascade that leads to spore germination^15,16^. Instead, *C. difficile* uses two soluble proteins located within the spore cortex, CspC and CspA. These two proteins are thought to sense germinant and co-germinant signals, respectively,^17,18^ and activate the CspB protease, which then proteolytically activates the cortex lytic enzyme SleC^19,20^. Activated SleC degrades the protective cortex, allowing germination to proceed^19^.

CspA, CspB, and CspC are members of the clostridial-specific subtilisin-like serine protease (Csp) family, which was originally identified in *Clostridium perfringens* as being responsible for the cleavage and activation of the cortex lytic enzyme SleC during germination^20,21^. Whereas the CspA, CspB, and CspC homologs encoded by *C. perfringens* and other Clostridiaceae and Lachnospiraceae family members are all predicted to be catalytically active, the only active Csp in *C. difficile* and other members of the Peptostreptococcaceae family is CspB^19,22,23^. *C. difficile* CspA and CspC harbor inactivating mutations in their catalytic triad and are thus pseudoproteases^19,22–24^. In *C. difficile*, the Csp proteins are encoded by the *cspBA-cspC* locus, with *cspBA* encoding a fusion of CspB and CspA (CspBA). During sporulation, CspBA undergoes interdomain processing by the YabG protease such that the individual CspB and CspA proteins are loaded into the mature spore^18,22^. Notably, CspA depends on CspC to be stably incorporated into the mature spore during sporulation^22,24^.

While prior genetic screens implicated CspC and CspA as the likely germinant and co-germinant receptors, respectively^17,18^, we previously showed that mutating residues within an unstructured region of CspC sensitizes *C. difficile* spores to both signals^25^. However, the mechanism by which CspC senses the germinant and co-germinant signals and the role of CspA in this process remained unclear. Here, we demonstrate that CspC and CspA form a stable heterodimer and determine its crystal structure. By mutating residues at the CspC:CspA interface, we identify specific regions that regulate the sensitivity of *C. difficile* spores to germinant and co-germinant signals. While we also show that CspA forms a homodimer and determine its crystal structure, mutating residues at the CspA homodimer interface minimally affect germinant and co-germinant signaling. Thus, the CspC:CspA heterodimer, rather than the individual CspC and CspA proteins, is a key signaling node for coordinately sensing germinant and co-germinant signals in *C. difficile*.

## Results

### CspC and CspA form a complex that does not include CspB

Previously, we showed that CspC levels in spores are greatly reduced in the absence of CspA^22,24^ and that CspC and CspA levels are also greatly reduced in the absence of CspB^24^. Therefore, we hypothesized that CspC and CspA directly interact and that this interaction may require CspB. To test these hypotheses, we co-expressed untagged CspA and CspB with His_6_-tagged CspC in *E. coli* and tested whether the untagged proteins co-purify with CspC-His_6_. CspA and CspB are produced as a CspBA fusion protein and then cleaved into individual proteins by the YabG protease during *C. difficile* sporulation^18,22^. To recapitulate the CspB and CspA variants generated by YabG-mediated processing of the CspBA fusion protein^18,22^, we produced the individual CspB_1-582_ and CspA_583-1132_ domains^18^. Since CspC and CspA have almost identical molecular weights (MW) of 60 kDa, we tagged CspC with a *Vibrio cholerae* MARTX toxin cysteine protease domain (CPD)^26^ fused to a C-terminal hexahistidine tag (CPD-His_6_, 24 kDa)^27,28^. CspB is readily distinguished from CspC and CspA because CspB undergoes autoprocessing of its N-terminal prodomain, and the processed CspB has a MW of ∼ 55 kDa^19^. The pull-down analyses revealed that untagged CspA, but not untagged CspB, efficiently copurifies with CspC-CPD-His_6_, suggesting that CspC and CspA form a complex (**Figure 1A**). CspA failed to co-purify with a GFP fusion protein (GFP-CPD-His_6_), used as a negative control, indicating that the CspC:CspA interaction is specific (**Figure 1A**). Interestingly, we found that CspC-CPD-His_6_ pulled down untagged CspBA fusion protein at considerably lower levels than untagged CspA (**Figure S1**), suggesting that YabG-mediated proteolytic processing of CspBA promotes CspC binding to CspA.

**Figure 1:**
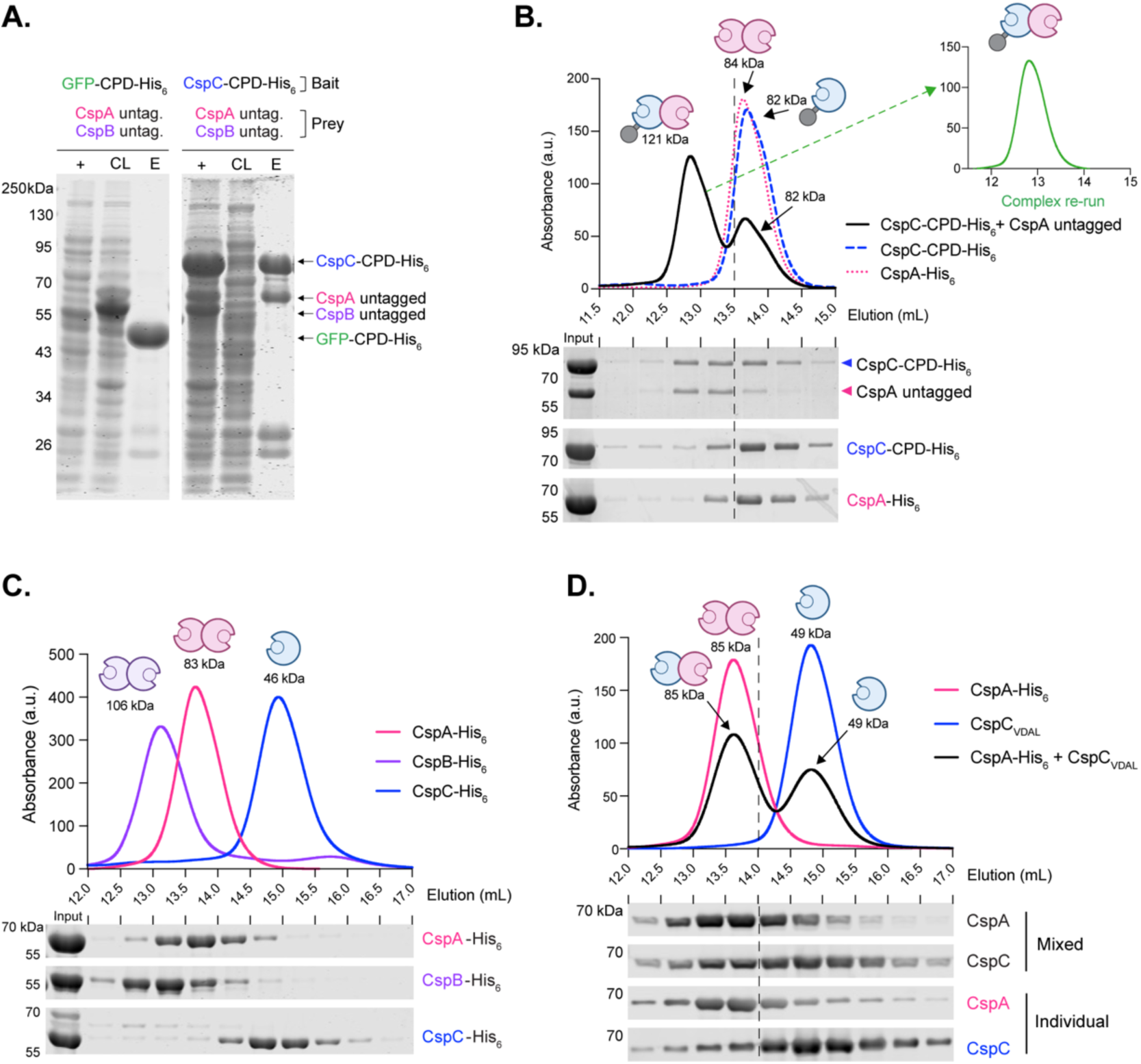
Interactions between *C. difficile* Csp proteins. (A) Coomassie stain of co-affinity purifications of CspC-CPD-His_6_ or GFP-CPD-His_6_ as the bait and untagged CspB and CspA as the prey. +, induced fraction; CL, cleared lysate; E, elution. (B) Size exclusion chromatography (SEC) analysis of the elution fraction of a CspC-CPD-His_6_:CspA co-affinity purification. The dashed line indicates the separation between the two peaks. 12.9 mL corresponds to an apparent MW of 121 kDa; 13.6 mL corresponds to an apparent MW of 84 kDa; and 13.7 mL corresponds to an apparent MW of 82 kDa. The SEC elution fractions were resolved using SDS-PAGE and stained with Coomassie. (inset) Analysis of the stability of SEC-purified CspC-CPD-His_6_:CspA complex. The complex was purified from the 12.6 – 13.0 mL fraction and then re-analyzed using SEC. (C) SEC analyses of individual Ni^2+^-affinity purified Csp-His_6_ proteins. 13.1 mL corresponds to an apparent MW of 106 kDa, 13.7 mL corresponds to an apparent MW of 83 kDa, and 14.9 mL corresponds to an apparent MW of 46 kDa. (bottom) Coomassie stain of SEC elution fractions resolved by SDS-PAGE. The CspB prodomain (∼9 kDa) is not shown. (D) Size exclusion chromatography (SEC) of separately purified CspC_VDAL_ and CspA-His_6_ variants. Separately purified proteins were mixed and incubated for 1 hr at a 1:1 stoichiometry and then resolved using SEC. (bottom) Western blot analysis of the SEC fractions. All data shown are representative of three replicates.

To assess the stability of the CspC-CPD-His_6_:CspA complex, we used size-exclusion chromatography (SEC) (**Figure 1B**). Two distinct peaks were observed by SEC. Peak 1 eluted with an apparent molecular weight (MW) of 121 kDa and contained CspA and CspC-CPD-His_6_ at an apparent ratio of 1:1 (**Figure 1B, SDS-PAGE**), consistent with a CspC-CPD-His_6_:CspA heterodimer. Notably, when the CspC-CPD-His_6_:CspA peak was re-run on SEC, it eluted as a single peak at the same volume as the original CspC-CPD-His_6_:CspA complex, indicating that this interaction is highly stable (**Figure 1B inset**). Peak 2 eluted with an apparent MW of 84 kDa. This peak contained mostly monomeric CspC-CPD-His_6_, consistent with its predicted MW of 86 kDa (**Figure 1B**). Despite CspA having a predicted MW of 60 kDa, some untagged CspA was detected in this smaller peak, suggesting that CspA may form a dimer. Consistent with this hypothesis, SEC analyses of individually purified CspC-CPD-His_6_ and CspA-His_6_ revealed that CspA-His_6_ forms an apparent homodimer, whereas CspC-CPD-His_6_ is monomeric, consistent with our previous study^25^ (**Figure 1C**).

To rule out any potential contributions of the CPD tag to this interaction, we individually purified CspC and CspA, mixed the two proteins at a 1:1 ratio, and analyzed the CspC:CspA heterodimer formation by SEC. Untagged CspC was first purified by inducing the CspC-CPD-His_6_ fusion protein to undergo autoprocessing using inositol hexakisphosphate (InsP_6_)^27,28^, which adds a Val-Asp-Ala-Leu (VDAL) sequence to the C terminus of CspC. When CspC_VDAL_ and CspA-His_6_ were mixed, a larger portion of the mixture eluted in peak 1 where the CspA homodimer typically elutes, consistent with CspC:CspA heterodimer formation (**Figure 1D**). These data indicate that CspA has a higher affinity for CspC than for itself and that the CspC:CspA heterodimer can form even when CspC and CspA are produced individually.

### CspB and CspA form homodimers, but CspC does not

Since CspA forms an apparent homodimer detected by SEC, we assessed the ability of *C. difficile* Csp proteins to interact in pairwise analyses. Untagged CspA, CspB, or CspC were co-produced with CPD-His_6_-tagged CspA, CspB, and CspC variants, respectively, in *E. coli*, and the resulting pairwise interactions were analyzed. As expected, untagged CspC co-purified with CspA-CPD-His_6_ (**Figure S2**), indicating that this interaction occurs when the tags are reversed. Untagged CspA co-purified with CspA-CPD-His_6_ (**Figure S2**), consistent with the ability of CspA to homodimerize. Untagged CspB also co-purified with CspB-CPD-His_6_ (**Figure S2**), suggesting that CspB forms a homodimer. Consistent with these data, SEC analyses of the individually purified Csp proteins (**Figure 1C**) confirmed that CspB and CspA each form homodimers. In contrast, untagged CspC was not pulled down by either CspB-CPD-His_6_ or CspC-CPD-His_6_ (**Figure S2**), and it eluted as a monomer when individually purified and analyzed via SEC (**Figures 1C**). None of the untagged Csps interacted with the GFP-CPD-His_6_ control (**Figure S2**). These results are consistent with the known crystal structures of the *C. perfringens* CspB dimer^16^ and *C. difficile* CspC monomer^20^ and provide the first evidence that *C. difficile* CspA forms a homodimer.

### CspA forms an antiparallel homodimer

To gain insight into the functional importance of the CspA homodimer uncovered by our co-affinity purification analyses, we determined the crystal structure of CspA. The structure was phased by molecular replacement using the known structure of the *C. difficile* CspC^25^ and refined to 3.2 Å resolution (**Table S3**). The *C. difficile* CspA structure is very similar to those of *C. difficile* CspC and *C. perfringens* CspB^19,25^ (**Figure 2A**) and can be superimposed with RMSDs of 1.22 and 1.44 Å, respectively. All three proteins are composed of an N-terminal prodomain followed by the subtilase domain that contains the central jellyroll domain insertion unique to the Csp subtilisin-like serine protease family^19,25^ (**Figure 2A & S3**).

**Figure 2:**
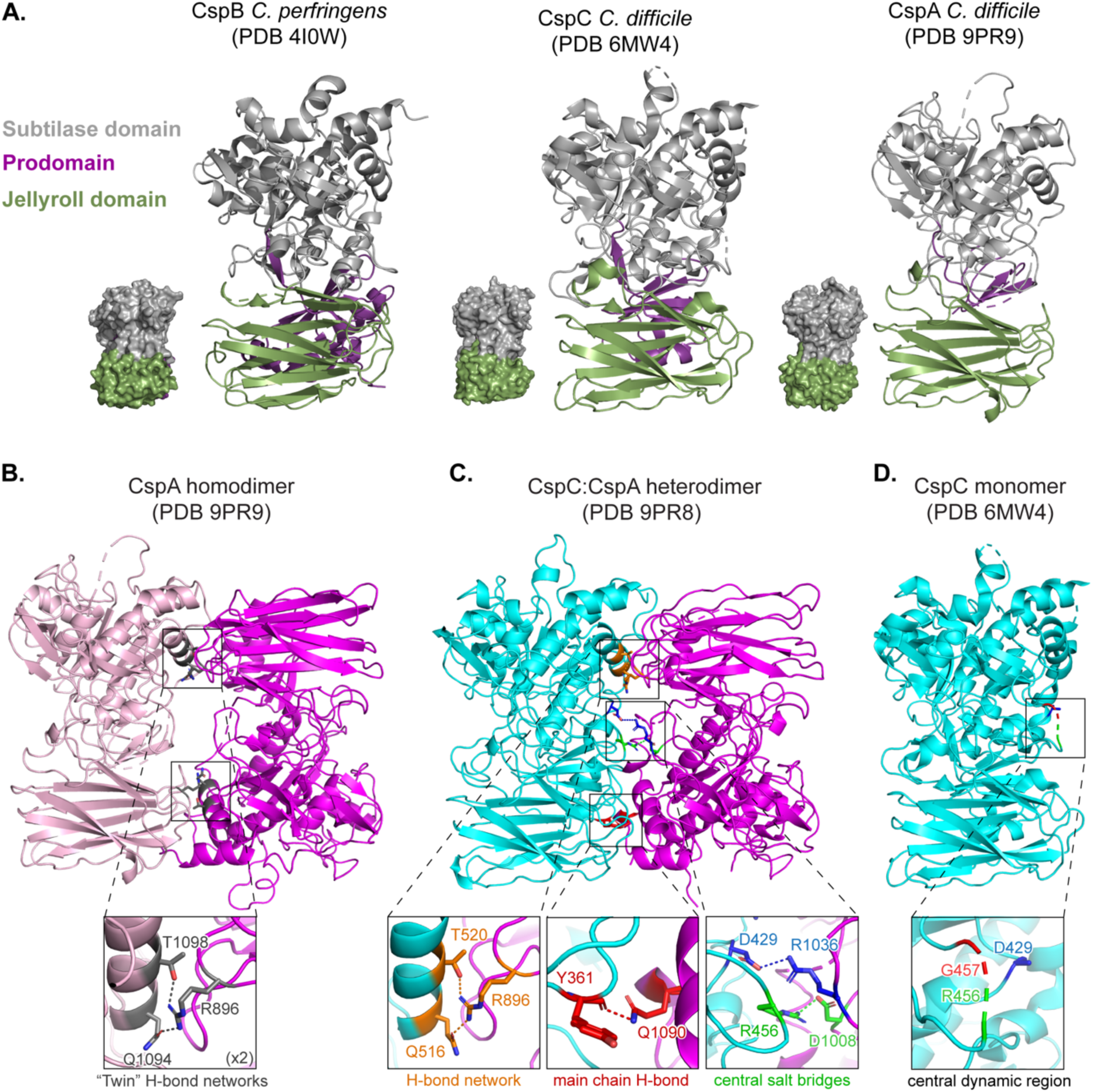
Crystal structures of the *C. difficile* CspA homodimer and CspC:CspA heterodimer. Ribbon and space-fill models of (A) *C. perfringens* CspB (PDB RI0W ^19^), *C. difficile* CspC (PDB 6MW4 ^25^), and *C. difficile* CspA (chain B) (this work – PDB 9PR9). The subtilase domains are shown in grey, jellyroll domains in green, and prodomains in purple. Ribbon models of the: (B) Crystal structure of the CspA homodimer solved to 3.2 Å (PDB 9PR9). Inset shows a close-up of one of two tripartite hydrogen bond networks found at the interface of the CspA homodimer. (C) Crystal structure of the CspC:CspA_F944E/Y1092E_ (CspC:CspA_EE_) heterodimer solved to 3.4 Å (PDB 9PR8). CspC is shown in cyan, and CspA is shown in magenta. Inset (left): tripartite hydrogen bond network between residues Q516_CspC_, T520_CspC_, and R896_CspA_ with distances of 2.7 Å and 3.1 Å, respectively. Inset (center): hydrogen bond between the Y361_CspC_ main chain and the Q1090_CspA_ residue with a distance of 2.8 Å. Inset (right): two salt bridges between CspC and CspA. The D429_CspC_, R1036_CspA_ salt bridge interaction is a distance of 3.6 Å, while the R456_CspC_, D1008_CspA_ is 2.9 Å. (D) Crystal structure of CspC monomer (PDB 6MW4 ^25^). Inset shows unstructured residues in the CspC monomer that form salt-bridges in the CspC:CspA heterodimer.

The asymmetric unit of the CspA crystals consists of two antiparallel, asymmetric homodimers, CspA_AB_ and CspA_CD_ (**Figure S4**). The four copies of CspA are very similar and can be superimposed with RMSDs ranging between 0.30 and 0.42 Å. Chains A and C are similar to each other and the most complete, with 534 residues resolved out of 559 total residues. Chains B and D are also similar to each other but less complete, with 489 and 487 residues, respectively, resolved out of 559 total residues. The two CspA homodimers can be superimposed with RMSD 0.32 Å. The N-terminal prodomains in chains A and C are better resolved than their counterparts in chains B and D in two regions **(Figure S4).** The A and C protomers are better resolved because of crystal contacts with neighboring protomers (**Figure S5**).

To map residues at the CspA homodimer interface, we analyzed the buried surface area and sidechain contacts (hydrogen bonds and salt bridges) using PiSA interface analysis^29^ (**Table S4**). In the CspA homodimer, the interface features two symmetrical tripartite hydrogen-bond (twin H-bond) networks formed between the jellyroll and subtilase domains (**Figure 2B**). Each H-bond network is formed by three residues (Arg896, Gln1094, and Thr1098; to maintain consistency, the numbering for CspA is presented based on the CspBA fusion protein) (**Figure 2B inset**). Notably, there were no interactions at the center of the homodimer (**Figure 2B**), so the twin H-bond networks make up a significant portion of the CspA homodimer interface.

### The CspC:CspA heterodimer has an extensive interface

To investigate the functional importance of the CspC:CspA heterodimer, we determined its crystal structure. Initial attempts to crystallize CspC_VDAL_:CspA failed to yield diffraction-quality crystals, so we hypothesized that the presence of the CspA homodimer in the CspC_VDAL_:CspA heterodimer preparations was interfering with the crystallization based on prior crystallization analyses (add protein from this reference), which can form a homodimer or a heterodimer with ()^30^. To mitigate this issue, we used a CspA_F944E/Y1092E_ (CspA_EE_) mutant that we had empirically determined to slightly increase CspC_VDAL_:CspA_EE_ heterodimer formation (**Figure S6**). This favoring of the heterodimer allowed for its crystallization^30^. The crystal structure of the CspC_VDAL_:CspA_EE_ heterodimer was determined to 3.35 Å resolution (**Figure 2C & Table S3**).

There are many similarities between the CspC:CspA heterodimer and the CspA homodimer. Like the CspA homodimer (**Figure 2B & S4**), the CspC:CspA heterodimer has an antiparallel orientation **(Figure 2C & Figure S7)**. Likewise, there are two crystallographically independent CspC:CspA heterodimers in the asymmetric unit, with small variations between their structures (RMSD of 0.40 Å between the two CspC:CspA heterodimers, 0.41 Å between CspC_B_ and CspC_D_, and 0.45 Å between CspA_A_ and CspA_C_) (**Figure S7**). In addition, the H-bond network formed by Arg896, Gln1094, and Thr1098 at the interface of the CspA homodimer, is conserved at the CspC:CspA heterodimeric interface. Specifically, the same residue, Arg896_CspA_, forms hydrogen bonds with Gln516_CspC_ and Thr520_CspC_, which are homologous to Gln1094_CspA_ and Thr1098_CspA_ (**Figure 2B inset & 2C left inset**).

However, there are also notable differences. First, the CspC:CspA heterodimer has a more extensive interface than the CspA homodimer (**Figures 2B & 2C**). Indeed, the heterodimeric interface of the CspC:CspA buries a significantly larger area, 2024 Å^2^ on average, compared to the homodimeric interface of CspA, which buries 1610 Å^2^ on average (**Table S4**). Second, in the CspC:CspA heterodimer, there is only one H-bond network at the interface (**Figure 2C**). Instead of the second H-bond network, a series of interactions tether the CspC jellyroll domain to the CspA subtilase domain, including a hydrogen bond between the main chain carbonyl group of CspC residue Tyr361 and the CspA residue Gln1090 side chain amide (**Figure 2C center inset**). Lastly, there are two salt bridges at the center of the interfaces of both heterodimers, between Asp429_CspC_ and Arg1036_CspA_ and between Arg456_CspC_ and Asp1008_CspA_ (**Figure 2C right inset**). Interestingly, the CspC residues forming these salt bridges are unstructured in the CspC monomer structure^25^ (**Figure 2C-D, insets**). Notably, the D chain Arg_CspC_ is unstructured in the CspC_D_:CspA_C_ heterodimer, so one of these salt bridges (Arg456_CspC_ and Asp1008_CspA_) is absent (**Figure S7C inset**). Previously, we showed that mutations of residues Asp429_CspC_, Arg456_CspC_, and Gly457_CspC_ enhance the sensitivity of spores to germinant and/or co-germinant signals^25^. These mutations would be expected to disrupt the salt bridges at the binding interface, potentially destabilizing the CspC:CspA heterodimer. Thus, the CspC:CspA heterodimer may be important for sensing both germinant and co-germinant signals.

### Mutation of the two salt bridges at the CspC:CspA heterodimer interface increases the sensitivity of *C. difficile* spores to TA germinant but has no effect on heterodimer stability

To dissect the functional roles of the aforementioned salt bridges, we mutated Asp429_CspC_, Arg456_CspC,_ Arg1036_CspA,_ and Asp1008_CspA_ (**Figure 2C & Figure 3A**) and tested the effect of the point mutations on *C. difficile* germination using an optical density (OD_600_)-based germination assay^17,18,25^ (**Figure S8**). The point mutations were introduced into *cspC* or *cspBA* complementation constructs expressed from an ectopic locus^31^ and the resulting mutants tested for their ability to rescue the germination of *ΔcspC* and Δ*cspBA* mutants.

**Figure 3.**
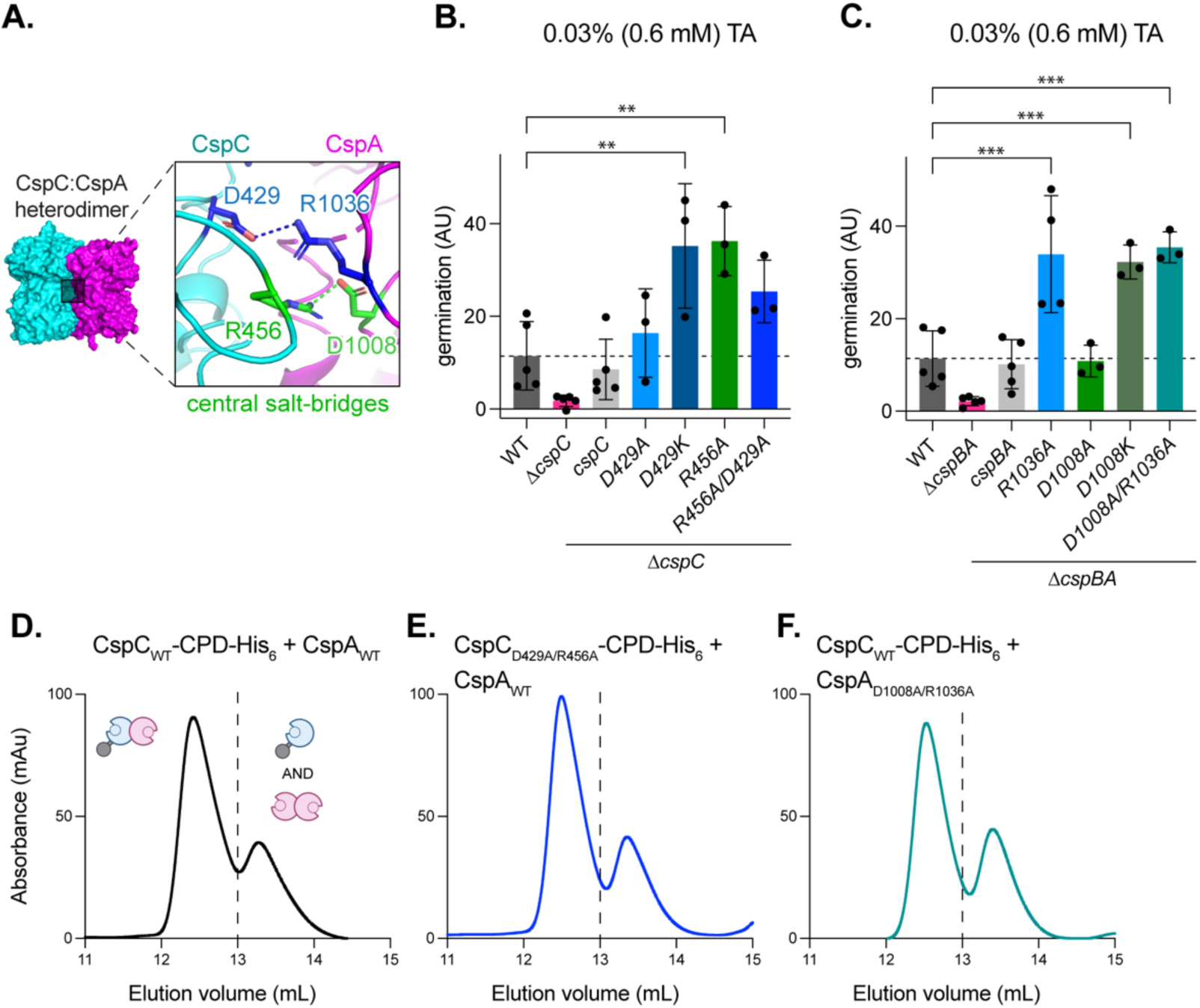
Mutation of two salt bridges at the CspC:CspA heterodimer interface increases the sensitivity of *C. difficile* spores to TA germinant. (A) Two salt bridges at the center of the CspC:CspA heterodimer interface. (B – C) Gemination levels based on the change in optical density (OD_600_) of purified spores suspended in rich medium over time following the addition of 0.6 mM taurocholate (TA). Germination in arbitrary units (AU) was calculated using the area below inverted OD_600_ curves (Figure S8). *cspC* complementation mutants were constructed in a *ΔcspC* background (B). *cspBA* complementation mutants were constructed in a *ΔcspBA* background (C). CspA residue numbers are based on the full-length CspBA fusion protein. Statistical significance relative to WT was determined using a one-way ANOVA and Dunnett’s multiple comparisons test. *** p < 0.001, ** p < 0.01. (D-F) Size exclusion chromatography analysis of CspC-CPD-His_6_ and CspA co-affinity purifications with the indicated mutants. The dashed line indicates the separation between the two peaks. All *C. difficile* data shown (B-C) are representative of a minimum of three independent replicates. The SEC data shown were obtained once.

Our prior work found that *cspC*_D429K_ spores were significantly more sensitive to TA germinant than the WT spores^25^. Here, we found that *cspC*_D429A_ spores were only slightly more sensitive to TA germinant, whereas *cspC*_R456A_ spores were significantly more sensitive to TA germinant (**Figure 3B**, p < 0.01). The reasons for differences in sensitivities to TA germinant are unclear, but they are consistent with our prior report showing that *cspC_R456G_* spores are more sensitive to TA germinant than *cspC*_D429K_ spores^25^. The double mutant *cspC*_D429A/R456A_ also produced spores with increased TA germinant sensitivity, albeit slightly less than spores carrying the single *cspC*_R456G_ mutation (**Figure 3B**).

*cspA*_R1036A_ and *cspA*_D1008K_ spores were significantly more sensitive to TA germinant (**Figure 3C**, p < 0.001), whereas *cspA*_D1008A_ spores exhibited WT TA germinant sensitivity (**Figure 3C**). *cspA*_D1008A/R1036A_ double mutant spores germinated similarly to the single *cspA*_R1036A_ mutant spores (**Figure 3C**). Western blot analyses confirmed that none of the salt-bridge substitutions altered Csp levels in spores relative to WT, although slight differences in mobility were observed with the CspC and CspA substitutions (**Figure S9**). These differences in mobility could be due to the altered charge of the Csp variants. Of note, increasing the resolution of the SDS-PAGE revealed multiple isoforms of CspA for the first time, which likely reflect differential processing of CspBA by YabG and an unknown protease^18,22^.

The location of these residues at the interface of the CspC:CspA heterodimer suggests that their substitution may destabilize the complex. To test the effect of these mutations on heterodimer stability *in vitro*, we used co-affinity purification followed by SEC analysis. Surprisingly, we found that substituting both the CspC and CspA residues involved in the two salt bridges for alanine (CspC_D429A/R456A_ or CspA_D1008A/R1036A_) had no effect on heterodimer stability *in vitro*, with both mutant heterodimers eluting like WT (**Figures 3E & 3F**). Taken together, these data show that mutations in the central salt bridge of the CspC:CspA heterodimer increase TA germinant sensitivity but have no effect on heterodimer stability *in vitro*.

### Mutations in the CspC:CspA heterodimer peripheral H-bonds reduce the sensitivity of *C. difficile* spores to TA germinant and have a minimal effect on heterodimer stability

To further probe the role of the CspC:CspA heterodimeric interface in *C. difficile* germination, we next targeted the H-bond network formed by the residues Gln516_CspC_, Thr520_CspC_, and Arg896_CspA_, using mutagenesis (**Figures 2E & 4A**). Previously, we showed that *cspC*_Q516E_ and *cspC*_Q516R_ spores had WT sensitivity to TA germinant^25^. Here, we found that single alanine substitutions *cspC*_Q516A_, *cspC*_T520A_, and *cspA*_R896A_ also had no effect on sensitivity to TA germinant at high concentrations (18.6 mM) (**Figure 4B-C**). However, double *cspC*_Q516A/T520A_ and triple *cspBA*_R896A_:*cspC*_Q516A/T520A_ mutant spores had decreased germinant sensitivity (**Figures 4B & D**). Moreover, substituting two or three H-bond network residues with glutamates (CspC_Q516E/T520E_ and CspBA_R896E_:CspC_Q516E/T520E_), which should disrupt the H-bond network by charge repulsion, completely abrogated germinant sensing (**Figures 4B & D**), and even the single glutamate substitution CspBA_R896E_ partially decreased germinant sensitivity (**Figure 4C**).

**Figure 4.**
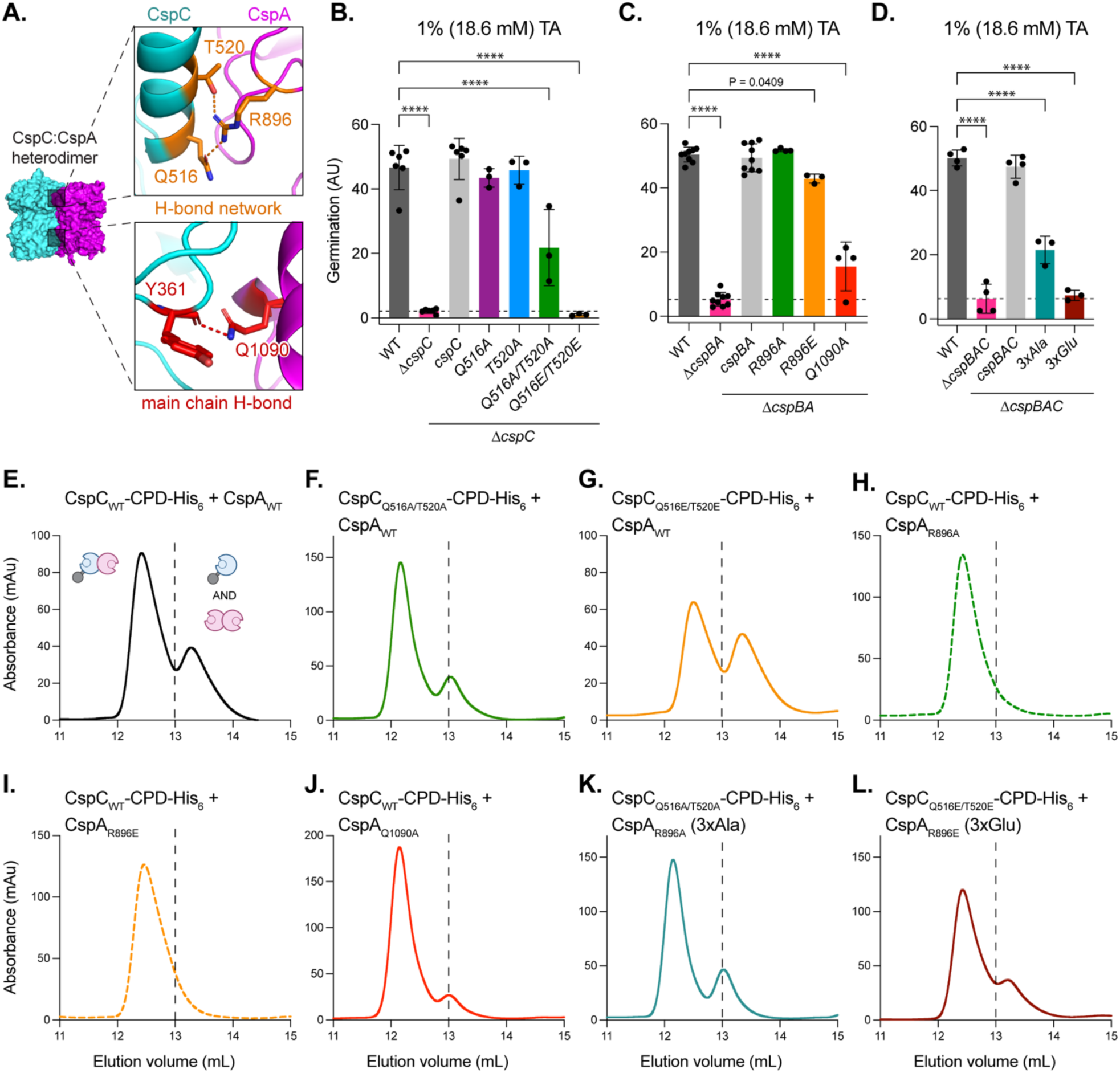
Mutations in the CspC:CspA heterodimer peripheral H-bonds impair germination and impact heterodimer formation. (A – top) Tripartite hydrogen bond network in the CspC:CspA heterodimer, and (A – bottom) hydrogen bond between the Y361_CspC_ main chain and the Q1090_CspA_ residue. (B, C) Germination levels based on the change in optical density (OD_600_) of purified spores suspended in rich medium over time following the addition of the 18.6 mM taurocholate (TA). Germination in arbitrary units (AU) was calculated using the area below inverted OD_600_ curves (Figure S8). *cspC* complementation mutants were constructed in a *ΔcspC* background (B). *cspBA* complementation mutants were constructed in a *ΔcspBA* background (C). *cspBAC* complementation mutants were constructed in a *ΔcspBAC* background (D). CspA residue numbers are based on the full-length CspBA fusion protein. *ΔcspBAC/3xAla* = triple-Ala substitution *cspBA_R896A_-cspC_Q516A/T520A_*, Δ*cspBAC/3xGlu* = triple-Glu substitution *cspBA_R896E_-cspC_Q516E/T520E_*. Statistical significance relative to WT was determined using a one-way ANOVA and Dunnett’s multiple comparisons test. **** p < 0.0001. (E-L) Size exclusion chromatography (SEC) analyses of CspC-CPD-His_6_:CspA co-affinity purifications. The dashed line indicates the separation between the two peaks. The trace for CspC_WT_-CPD-His_6_:CspA_WT_ is reproduced from Figure 3D. All *C. difficile* data shown (B-D) are representative of a minimum of three independent replicates. E-L are representative of a minimum of two independent replicates, except for CspC_WT_-CPD-His_6_:CspA_Q1090A_, which was resolved via SEC once.

We next probed the role of a single hydrogen bond between the main chain carbonyl group of residue Tyr361_CspC_ and residue Gln1090_CspA_ on the reciprocal side of the heterodimer interface from the H-bond network (**Figure 2C & 4A**) by substituting residue Gln1090_CspA_ for alanine. Similarly to the heterodimer H-bond network mutations, the *cspA*_Q1090A_ mutation greatly reduced TA germinant sensing even at high concentrations (18.6 mM) (**Figure 4C**, p < 0.0001). None of the mutations affected Csp levels in spores (**Figure S10**), indicating that the observed germination defects are not due to reduced levels of germination proteins in spores. Taken together, these data suggest that disrupting the H-bonds at the periphery of the CspC:CspA heterodimer interface impairs TA germinant sensing.

To test the effects of peripheral H-bond substitutions on heterodimer stability *in vitro,* we used co-affinity purification followed by SEC (**Figure 4E - L**). Most mutations had no effect on the CspC:CspA heterodimer stability. However, the double glutamate mutant CspC_Q516E/T520E_ reduced CspC:CspA heterodimer stability (**Figure 4G**) and even decreased the amount of untagged CspA pulled down by CspC-CPD-His_6_ during the co-affinity purification (**Figure S11A**). The CspC_Q516E/T520E_ mutant is one of two germination-null mutants (**Figure 4B**), which suggests that the disruption of the H-bond network in the CspC:CspA heterodimer can reduce both the heterodimer stability and TA germinant sensing.

All other mutants tested eluted similarly to the WT heterodimer, apart from the CspA_R896A_ and CspA_R896E_ variants, which eluted exclusively in complex with CspC (**Figure 4H-I**). Notably, Arg896_CspA_ also participates in H-bond networks at the CspA homodimer interface (**Figure 2B inset**). Since the second peak in the WT CspC-CPD-His_6_:CspA SEC trace contains both the CspC monomer and some CspA homodimer (**Figure 1B and 4E**), disrupting the interactions formed by CspA Arg896 likely impairs CspA homodimerization, promoting CspC:CspA interaction.

Like the CspC_Q516E/T520E_ mutant, the triple glutamate substitution in the H-bond network of the CspC:CspA heterodimer (CspC_Q516E/T520E_:CspA_R896E_) abrogated germination at high concentrations of TA germinant (**Figure 4D**). However, unlike the double glutamate mutant CspC_Q516E/T520E_ (**Figure 4G**), the triple mutant CspC_Q516E/T520E_:CspA_R896E_ eluted like the WT heterodimer (**Figure 4L**). Since CspA_R896E_ may impair CspA homodimerization, promoting CspC:CspA interaction, it is possible that combining this mutant with CspC_Q516E/T520E_ counterbalances the destabilizing effect of the CspC mutations, resulting in an SEC trace similar to the WT heterodimer. Regardless, these data reveal that, while there is no direct correlation between heterodimer stability *in vitro* and TA germinant sensitivity in spores, the H-bond network at the interface of CspC:CspA is important for heterodimerization and critical for germinant sensing.

### Mutations in the CspA H-bond networks disrupt CspA homodimerization but have variable effects on germination

While our analyses pointed to the importance of the CspC:CspA heterodimer in *C. difficile* germinant signaling, the role of the CspA homodimer detected *in vitro* (**Figures 1 & 2**) in germination remained unclear. To address this, we designed mutations targeting the twin H-bond networks in the CspA homodimer, formed by Arg896, Gln1094, and Thr1098 (**Figure 2B & 5A**). Arg896_CspA_ is also involved in the analogous H-bond network in the CspC:CspA heterodimer **(Figure 2B inset & C left inset)**, and Arg896_CspA_ mutants reduced *C. difficile* TA germinant sensing and favored the CspC:CspA heterodimer formation *in vitro* (**Figure 4**). To further interrogate the role of the twin H-bond networks in the CspA homodimer, we generated *cspA* mutants encoding double and triple substitutions targeting H-bond network residues. Similarly to *cspBA*_R896E_ mutant spores, *cspBA*_Q1094E/T1098E_ mutant spores were less sensitive to TA germinant at both low and high germinant concentrations (**Figure 5B & C**). While the *cspBA*_Q1094E/T1098E_ mutant spores were less sensitive to TA germinant at high concentrations of germinant (18.6 mM, **Figure 5C**), these phenotypes were relatively subtle compared to the analogous mutations in the CspC:CspA heterodimer H-bond network, which completely abrogated TA germinant sensing (*cspC*_Q516E/T520E_, **Figure 4B**). By contrast, *cspBA*_R896E/Q1094E_ and *cspBA*_R896A/Q1094A/T1098A_ (*3xAla*) mutant spores were more sensitive to low concentrations of TA germinant (0.6 mM) (**Figure 5B**). Since none of the CspA mutations impacted Csp levels in spores in western blot analyses (**Figure S12**), these data indicate that mutation of the CspA homodimer H-bond networks had relatively minor and variable effects on germinant sensitivity compared to the analogous mutations of the H-bond network in the CspC:CspA heterodimer (**Figure 4**).

**Figure 5.**
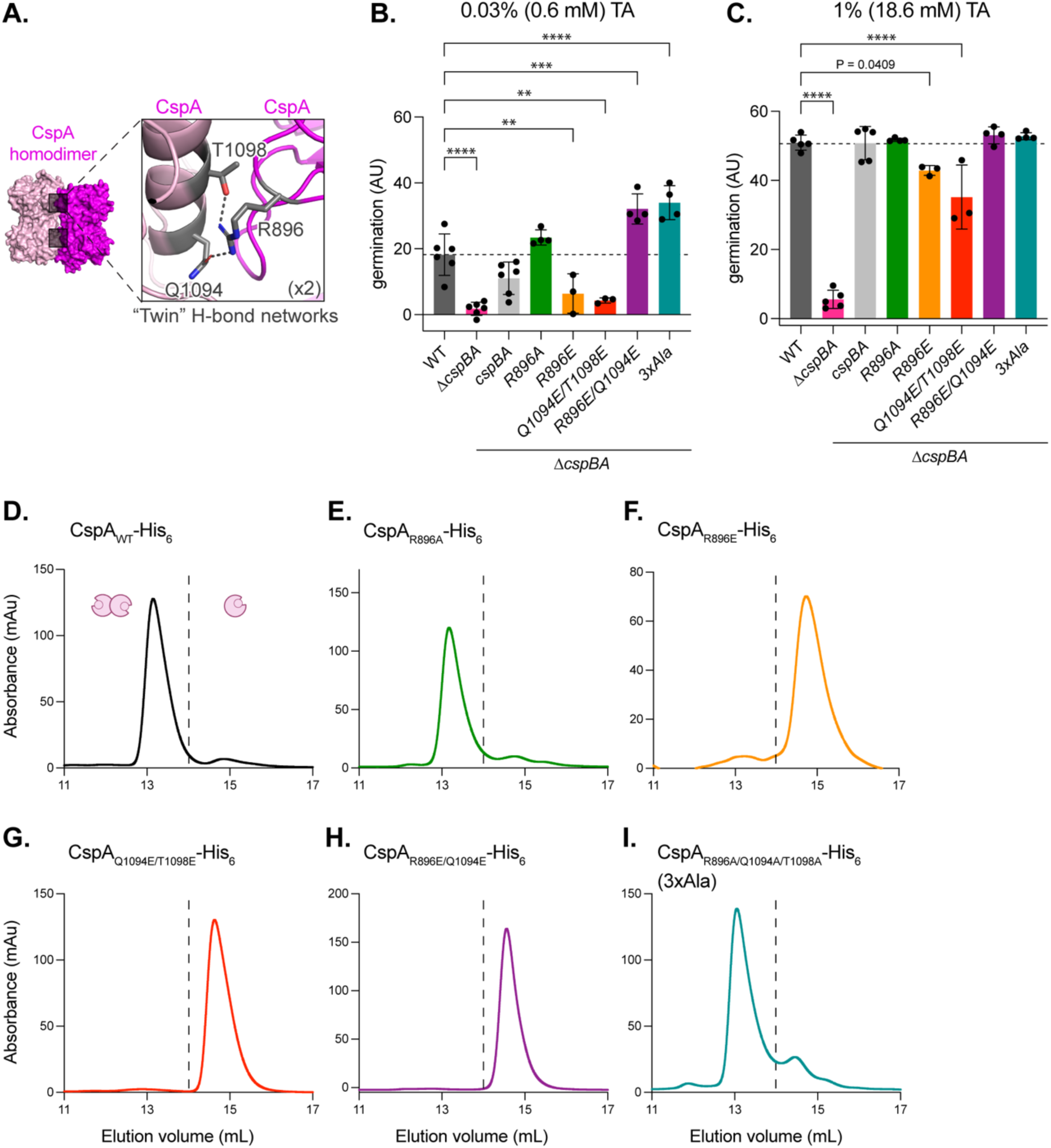
Mutations in the CspA twin H-bond networks disrupt CspA homodimerization but have variable effects on germination. (A) One of the two tripartite hydrogen bond networks at the CspA homodimer interface. (B-C) Germination levels based on the change in optical density (OD_600_) of purified spores suspended in rich medium over time following the addition of the indicated concentration of taurocholate (TA). Germination in arbitrary units (AU) was calculated using the area below inverted OD_600_ curves (Figure S8). *cspBA* complementation mutants were constructed in a *ΔcspBA* background. CspA residue numbers are based on the full-length CspBA fusion protein. Δ*cspBA/3xAla* = *cspBA_R896A/Q1094A/T1098A_*. The *cspBA_R896A_* and *cspBA_R896E_* measurements presented in (C) are reproduced from Figure 4C. Statistical significance relative to WT was determined using a one-way ANOVA and Dunnett’s multiple comparisons test. **** p < 0.0001, *** p < 0.001, ** p < 0.01. (D-I) Size exclusion chromatography (SEC) analyses of CspA-His_6_ variants following affinity purification. The dashed line indicates the separation between the two peaks. All *C. difficile* data shown (B-C) are representative of a minimum of three independent replicates. The data shown in D-I were obtained once, except for CspA_R896A_-His_6_, which was resolved via SEC using three independent replicates.

To determine how the H-bond network substitutions affected CspA homodimerization *in vitro*, we analyzed the SEC profiles of purified CspA-His_6_ variants. While alanine substitutions in H-bond network residues (CspA_R896A_-His_6_ and CspA_R896A/Q1094A/T1098A_-His_6_) did not affect CspA homodimerization (**Figure 5E & I**), even single glutamate substitutions were sufficient to disrupt the CspA homodimer formation (CspA_R896E_-His_6_, CspA_Q1094E/T1098E_-His_6_, and CspA_R896E/Q1094E_-His_6_, **Figure 5F-H**). Importantly, we found no correlation between the effect of the mutations on CspA homodimer stability and *C. difficile* spore sensitivity to TA germinant. The double glutamate substitution *cspBA*_R896E/Q1094E_ disrupted CspA homodimerization *in vitro* (**Figure 5H**) and slightly increased spore sensitivity to TA germinant (**Figure 5B**), whereas other glutamate substitutions (CspA_R896E_ and CspA_Q1094E/T1098E_) disrupted CspA homodimer formation (**Figure 5F & G**) and decreased spore sensitivity to TA germinant (**Figure 5B & C**). Further, CspA substitutions in the other residues of the homodimer H-bond network stabilized the CspC:CspA heterodimer *in vitro* in SEC analyses **(Figure S13)**, further supporting the hypothesis that disruption of the CspA homodimer promotes the CspC:CspA heterodimer formation. Since no notable correlation between CspA homodimer formation *in vitro* and *C. difficile* spore TA germinant sensitivity was observed, our data suggest that CspA homodimerization, which occurs under *in vitro* conditions, has little, if any, biological significance.

### The CspC:CspA heterodimer is extremely stable in the presence of germinants and co-germinants

Since our mutagenesis revealed that residues at the CspC:CspA heterodimer interface are important for germinant sensing (**Figures 3-4**), we considered the possibility that disruption of the CspC:CspA heterodimer is a key step in the TA germinant (and, perhaps, even co-germinant) sensing. However, despite extensive mutagenesis across the CspC:CspA heterodimer interface, we identified only a single CspC variant that partially disrupted the heterodimer *in vitro* (CspC_Q516E/T520E_, **Figure 4G**). This finding strongly suggests that the CspC:CspA heterodimer is extremely stable. To assess its stability, we performed limited proteolysis on the CspC-CPD-His_6_:CspA heterodimer, the CspC-CPD-His_6_ monomer, and the CspA-His_6_ homodimer (**Figure 6A**). Consistent with prior work^25^, the CspC monomer was sensitive to high levels of chymotrypsin (**Figure 6A**). The CspA homodimer was similarly sensitive to chymotrypsin (**Figure 6A**). By contrast, the CspC:CspA heterodimer was highly resistant to proteolysis by chymotrypsin.

**Figure 6.**
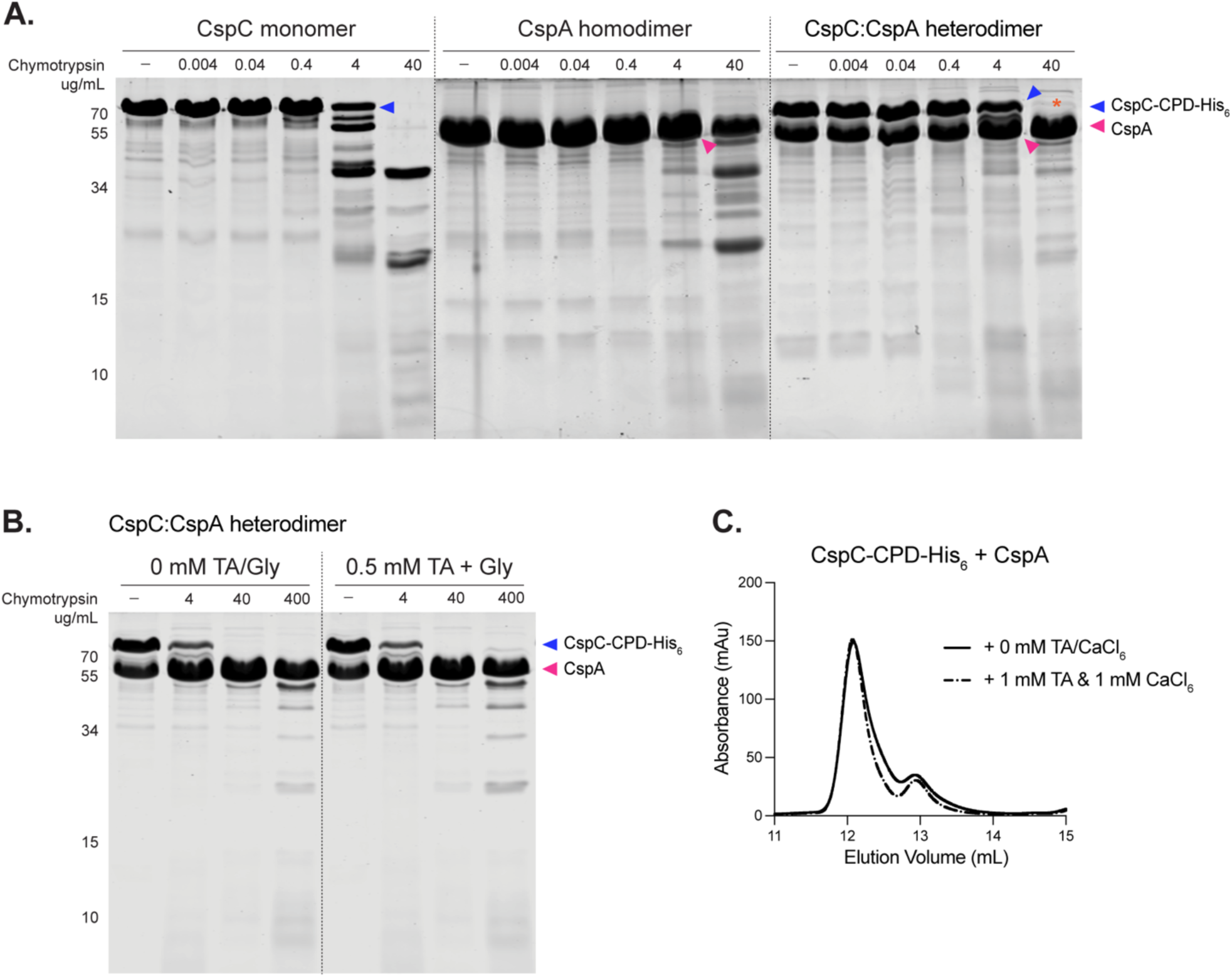
The CspC:CspA heterodimer is extremely stable in the presence and absence of germinants and co-germinants. (A-B) Limited proteolysis analyses of purified CspC, CspA, and the CspC-CPD-His_6_:CspA heterodimer using the indicated concentrations of chymotrypsin. CspA-His_6_ and CspC-CPD-His_6_:CspA heterodimer were affinity-purified. The CspC monomer was affinity-purified as a CspC-CPD-His_6_ fusion and then the CPD-His_6_ tag was removed by InsP_6_ addition. Cleavage of the CspC-CPD-His_6_:CspA heterodimer at 40 μg/mL chymotrypsin is most likely due to digestion of the CPD-His_6_ tag. (B) Limited proteolysis of purified CspC-CPD-His_6_:CspA heterodimer in the presence and absence of the indicated concentrations of taurocholate (TA) and glycine (Gly). Ca^2+^ co-germinant was not tested in this assay because high concentrations of CaCl_2_ inhibit chymotrypsin activity. (C) Size exclusion chromatography analyses of CspC-CPD-His_6_:CspA co-affinity purifications in the presence or absence of the indicated concentrations of TA and CaCl_2_. All data shown are representative of two independent replicates, except for C, which was obtained once.

We next considered the possibility that germinants and co-germinants may disrupt the heterodimer or induce conformational changes that might be detectable by limited proteolysis. However, dual exposure to TA germinant and the co-germinant glycine did not alter the protease sensitivity of the CspC:CspA heterodimer (**Figure 6B**). We next assessed whether the CspC-CPD-His_6_:CspA heterodimer might be destabilized during SEC in the presence of the TA germinant and the co-germinant calcium, since this method does not analyze proteins at equilibrium. No difference in the SEC profiles of the heterodimer was observed in the presence or absence of germinant and co-germinant (**Figure 6C**). Thus, the presence of the TA germinant and the co-germinants does not appear to affect the stability or overall conformation of the CspC:CspA heterodimer *in vitro*.

### Two regions of the CspC:CspA heterodimer function independently to regulate (co)germinant signal integration

Our mutational analyses revealed that disrupting interactions across two distinct regions of the heterodimer interface results in markedly distinct phenotypes: the salt bridge substitutions enhance germinant sensitivity^25^ (**Figure 3**), while the peripheral H-bond substitutions impair germination (**Figure 4**). However, it remained unclear whether each region plays a functionally distinct, independent role in signaling or whether the regions signal through a linear, i.e., hierarchical, pathway. To distinguish between these possibilities, we combined mutations with the strongest germination phenotypes in a quadruple mutant *ΔcspBAC/cspBA*_D1008A/R1036A_:*cspC*_Q516E/T520E_ (*4x mut*) (**Figure 2C & 6A**). We then used this mutant to identify whether one region functionally constrains the other – in other words, is epistatically dominant – or if the two regions function independently, in parallel^32,33^.

If one region exhibits epistasis over the other, then a mutant containing substitutions in both regions would have the same phenotype as the mutant containing substitutions only in the dominant region^32^. Instead, we found that 4x mutant spores exhibited an intermediate germinant phenotype: they germinated better than the germination-null *cspC*_Q516E/T520E_ mutant spores but worse than the hypersensitive *cspA*_D1008A/R1036A_ mutant spores (**Figure 7B-C**). Notably, the 4x mutations resulted in slightly decreased levels of Csps in dormant spores (∼ 30%, **Figure S14**). While this modest decrease in Csp levels could partially contribute to reduced germination, the *cspC*_Q516E/T520E_ and *cspA*_D1008A/R1036A_ mutations had no impact on Csp levels in spores (**Figures S9B & S10A**), and their phenotypes reflect changes in CspC:CspA function rather than abundance. Therefore, the 4x mutant phenotype likely reflects a partial contribution from both regions, suggesting that neither region is epistatically dominant over the other. Thus, signal transduction through these regions of the CspC:CspA heterodimer does not occur in a hierarchical fashion and, instead, the two regions likely signal through independent parallel pathways.

**Figure 7.**
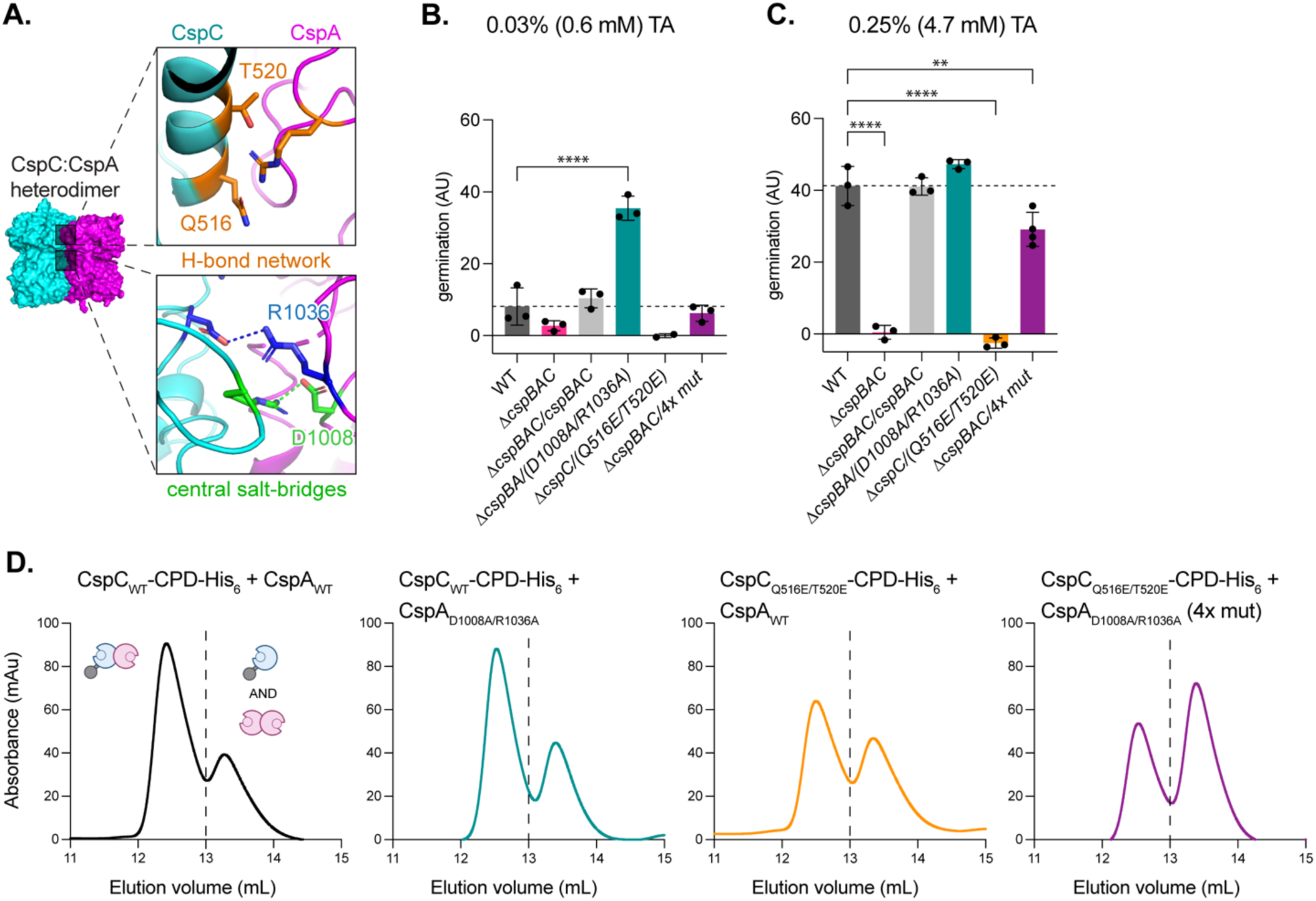
Two regions of the CspC:CspA heterodimer function independently to regulate germinant and co-germinant signal integration. (A – top) Tripartite hydrogen bond network in the CspC:CspA heterodimer, and (A – bottom) salt bridges at the center of the CspC:CspA heterodimer interface. (B-C) Gemination levels based on the change in optical density (OD_600_) of purified spores suspended* in rich medium over time following the addition of 0.6 mM taurocholate (TA) (B) or 4.7 mM TA (C). Germination in arbitrary units (AU) was calculated using the area below inverted OD_600_ curves (Figure S8). Germination phenotypes of double and quadruple mutants were best resolved at 4.7 mM TA (Figure S8). *cspBAC* complementation mutants were constructed in a *ΔcspBAC* background; *cspBA* complementation mutant was constructed in a *ΔcspBA* background; and *cspC* complementation mutants were constructed in a *ΔcspC* background. CspA residue numbers are based on the full-length CspBA fusion protein. Δ*cspBAC/4x mut* = *cspBA_D1008A/R1036A_-cspC_Q516E/T520E_*. Statistical significance relative to WT was determined using a one-way ANOVA and Dunnett’s multiple comparisons test. **** p < 0.0001, ** p < 0.01. (D) Size exclusion chromatography (SEC) analyses of CspC-CPD-His_6_:CspA co-affinity purifications. The dashed line indicates the separation between the two peaks. The traces for CspC_WT_-CPD-His_6_:CspA_WT_, CspC_WT_-CPD-His_6_:CspA_D1008A/R1036A_, and CspC_Q516E/T520E_-CPD-His_6_:CspA_WT_ are reproduced from Figures 3D, 3F, and 4G, respectively. All *C. difficile* data shown (B-C) are representative of a minimum of three independent replicates. The data shown in D were obtained twice, except for CspC-CPD-His_6_:CspA_D1008A/R1036A_, which was resolved via SEC once.

We next analyzed the effect of the 4x substitutions on the stability of the CspC:CspA heterodimer using recombinant proteins. We found that combining the 4x substitutions decreased CspC:CspA heterodimer stability even more than the CspC_Q516E/T520E_ variant, with less untagged CspA_D1008A/R1036A_ being pulled down by the CspC_Q516E/T520E_-CPD-His_6_ variant during affinity purification compared with WT and the constituent CspA or CspC mutants alone (**Figure S11B**). Furthermore, the 4x mutant heterodimer was less stable in SEC than the CspC_Q516E/T520E_ mutant (**Figure 7D**). These data Notably, despite targeting two regions of the CspC:CspA heterodimer, the 4x mutant did not fully disrupt heterodimer formation *in vitro* (**Figure 7D**). This finding suggests that the Y361_CspC_:Q1090_CspA_ main chain H-bond contributes to the stability of the CspC:CspA heterodimer (**Figure 2C, middle inset & 4A, lower inset)**. Importantly, once again, we found no direct correlation between germinant sensitivity and CspC:CspA heterodimer stability *in vitro* (**Figure 7B-D**). Taken together, our data suggest that while the CspC:CspA heterodimer plays a central role in germinant sensing, its stability alone does not determine germinant sensitivity. These data also support the idea that different regions at the heterodimeric interface function independently to mediate germinant signal transduction.

## Discussion

Genetic screens originally implicated the CspC and CspA pseudoenzymes as the germinant and co-germinant receptors in *C. difficile*^17,18^, respectively. However, our subsequent structure-function analyses suggested that CspC is involved in sensing both signals during germination^25^, so it was unclear how CspC cooperates with CspA to regulate both germinant and co-germinant sensing. Here, we address this question by demonstrating that CspC and CspA form a highly stable heterodimer (**Figure 8A**). Our extensive mutational analysis of the CspC:CspA heterodimeric interface, informed by the crystal structure of the heterodimer determined here, revealed that the CspC:CspA heterodimer plays a key role in sensing germinant and co-germinant signals in *C. difficile*. Specifically, we discover key interaction sites that regulate *C. difficile* germinant sensing: a central dynamic region that controls the sensitivity of *C. difficile* spores to germinant signals and peripheral interactions that are critical for germinant signal transduction. Moreover, our structure-function analyses identify, for the first time, CspA residues that regulate the sensitivity of *C. difficile* spores to TA germinant (**Figures 3-6**). Thus, both CspC and CspA, rather than the individual proteins^25^, determine the sensitivity of *C. difficile* spores to *both* germinant and co-germinant signals because it is the CspC:CspA complex, rather than the individual proteins, that integrates and transduces germinant and co-germinant signals in *C. difficile*.

**Figure 8.**
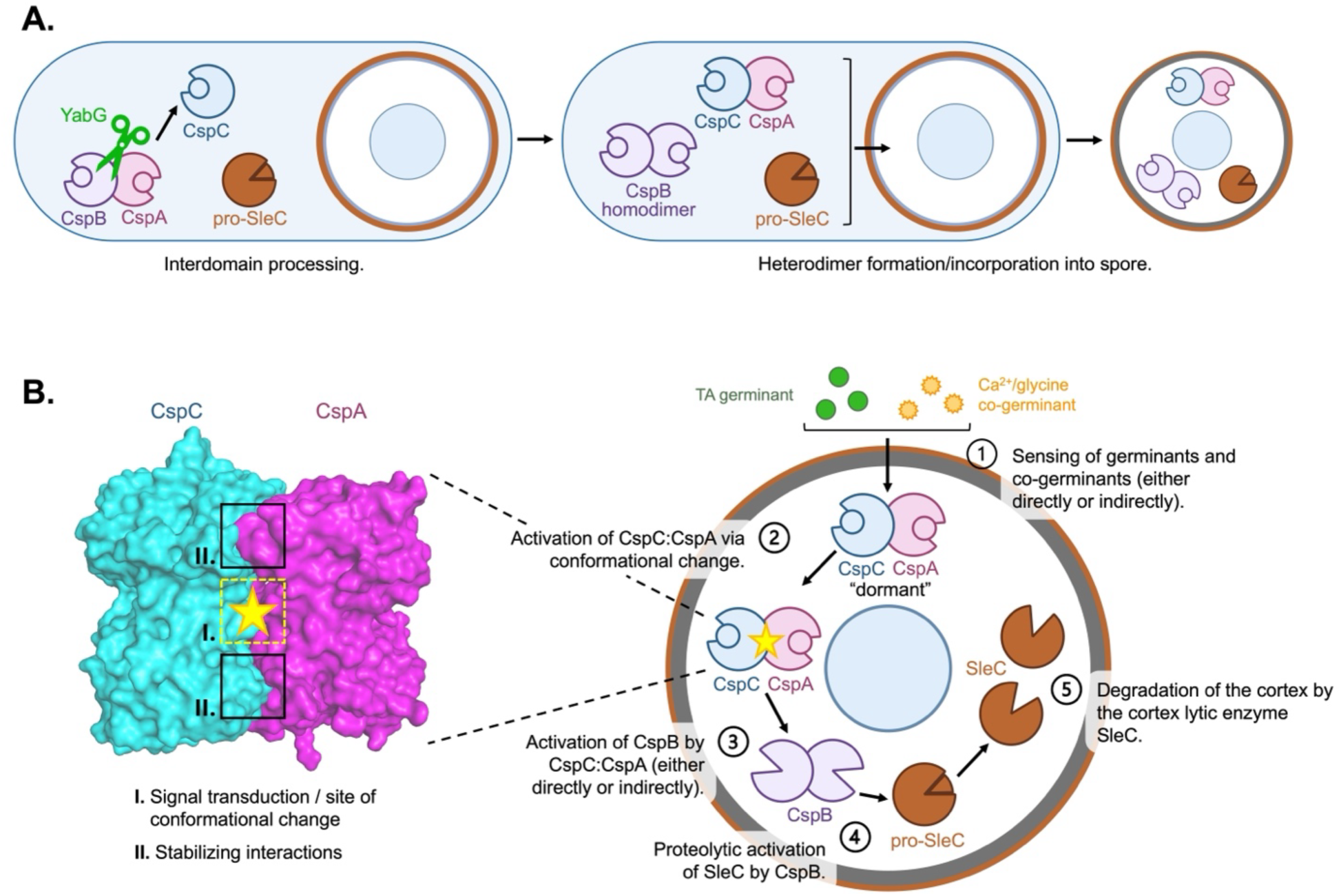
Model for *C. difficile* germination. (A) Model of the *C. difficile* proteins required for germination during sporulation. From left to right: interdomain processing of the CspBA fusion protein by the protease YabG; heterodimerization of CspC and CspA and homodimerization of CspB; and loading of key germination proteins into the cortex region of the mature spore ^54^. (B) Model of a *C. difficile* spore undergoing germination. Sensing of germinant and co-germinant signals (either directly or indirectly) by the CspC:CspA heterodimer (1) leads to the heterodimer adopting an active conformation (2). Inset: CspC:CspA heterodimer space-filling model. Residues at the center of the CspC:CspA heterodimer binding interface (I.) play important roles in transducing germinant and co-germinant signals, while residues at the periphery of the binding interface (II.) help stabilize the complex. Signal transduction by the CspC:CspA heterodimer initiates the activation of CspB (either directly or indirectly) (3). Active CspB then proteolytically activates the cortex-lytic enzyme SleC (4), which will go on to degrade the spore cortex (5), allowing for rehydration of the spore core and resumption of metabolic activity.

While we also demonstrate for the first time that CspA forms a homodimer *in vitro*, our structure-function analyses, guided by our newly determined crystal structure, strongly suggest that disrupting CspA homodimerization has only subtle effects on *C. difficile* TA germinant sensing compared to analogous mutations in the CspC:CspA heterodimer (**Figures 4-5**). Therefore, we propose that CspA homodimerization is less physiologically relevant than CspC:CspA heterodimerization during *C. difficile* spore germination.

Our interrogation of the CspC:CspA binding interface has yielded key insights into the mechanism by which the heterodimer transduces signals during *C. difficile* spore germination. While we previously showed that the CspC residues, Asp429, Arg456, and Gly457 – which are unstructured in the CspC monomer – influence the sensitivity of spores to both germinant and co-germinant signals^25^, we now show that these residues are part of a central dynamic region of the heterodimer. In one heterodimer, these residues form two salt bridges with CspA (**Figures 2C & 3A**), while in the other heterodimer of the asymmetric unit, they form a single salt bridge (**Figure S7**). Similar to our prior analyses of the CspC residues that participate in these salt-bridge interactions^25^, we found that mutation of the CspA residues engaged in these salt-bridge interactions hypersensitize spores to TA germinant (**Figure 3**). This suggests that the central salt bridges in the CspC:CspA heterodimer normally inhibit germinant and co-germinant signaling in dormant spores. Moreover, the conformationally dynamic nature of this region gleaned from the crystal structures reported here and previously^25^ suggests that this region undergoes conformational changes during germinant and co-germinant sensing to facilitate signal transduction (**Figure 8B**). We hypothesize that the two heterodimer structures reported here may represent “dormant” vs. "active", or "primed", conformational states (**Figures S7 & 8B**) that regulate *C. difficile* germination.

By contrast, the H-bonds at the periphery of the heterodimeric interface appear to be less conformationally dynamic (**Figure 2 & Table S4**). Their disruption abrogates germinant sensing and reduces heterodimer formation (**Figure 4**), which suggests that heterodimerization is important for germinant sensing and subsequent signal transduction. Interestingly, we found that the central dynamic region and the peripheral H-bonds function independently during germinant signal transduction (**Figure 7B**), with neither region exhibiting protein epistasis over the other. These findings suggest a model where the peripheral H-bonds stabilize the pseudoproteases in the heterodimeric state, which is required for the central dynamic region to transduce the germination signal (**Figure 8B**). Signal transduction through this central region by the CspC:CspA heterodimer then initiates the activation of the CspB protease, which in turn proteolytically activates the cortex lytic enzyme SleC^19,20^ (**Figure 8**), triggering cortex degradation and the resumption of metabolic activity^5,8^.

Our discovery of the CspC:CspA heterodimer also suggests an explanation for why CspC requires CspA for its stable incorporation into spores^22,24^. Given the extensive interactions between CspC and CspA revealed by the CspC:CspA crystal structure, we hypothesize that CspA binding to CspC is necessary for CspC to be stably incorporated into spores. Since we were unable to identify mutants that fully disrupt heterodimer formation, it was not possible to test this hypothesis, but our finding that Csp levels are slightly reduced (∼30%) in the 4x mutant (**Figure S14**) is consistent with this hypothesis, since these four mutations decrease CspC:CspA heterodimer formation *in vitro* (**Figure 7**).

Regardless, with the identification of the CspC:CspA heterodimer as a key signaling node, our findings raise three key questions: (1) Does the CspC:CspA heterodimer directly sense germinants and co-germinants? (2) What conformational changes occur in the CspC:CspA heterodimer in response to the germinant and co-germinant sensing? 3) How does the CspC:CspA heterodimer activate its cognate protease, CspB? Studies of pseudoenzyme function in diverse systems provide a useful framework for considering these questions because pseudoenzymes, like CspC and CspA, often modulate the activity of their cognate enzymes using a variety of mechanisms^34–36^.

As for the first question, pseudoenzymes often act as molecular switches that bind small-molecule ligands, which alter their conformation and allow them to activate their cognate enzymes^36^. According to this model, germinant and co-germinant binding to the CspC:CspA heterodimer would cause the conformational changes that trigger germination. While we were unable to detect any effect of the germinants and co-germinants on the CspC:CspA heterodimer formation or stability *in vitro* (**Figure 6**), these assays did not measure binding directly. However, detecting direct binding between germinants and their receptors has been challenging even in *B. subtilis*, despite strong genetic evidence that germinants are recognized by nutrient-gated channel-like Ger receptors^15,37,38^. Moreover, these *in vitro* assays could be missing key regulatory factors found within *C. difficile* spores or the environmental signals necessary for these interactions. For example, the gel-forming GerG protein regulates Csp levels in spores and spore germinant sensitivity^39^, so proper CspC:CspA function may depend on GerG. Indeed, the absence of regulatory proteins like GerG could explain why our *in vitro* analyses of CspC:CspA heterodimer stability do not correlate strongly with our germination phenotypes (**Figure 7**). We aim to address the involvement of other important proteins, like GerG, during *C. difficile* germination in future studies.

For the second question, our analyses identified key residues in the heterodimer that are likely involved in conformational changes that drive germination, and our crystal structures reveal several conformationally dynamic regions in both the CspC:CspA heterodimer and the CspA homodimer that may contribute to this pathway. Notably, the N-terminal prodomain of CspA is partially unstructured in two of the chains in the CspA homodimer asymmetric unit, but structured in the other two chains (**Figure S4**). Interestingly, CspA’s N-terminal prodomain is generated when the YabG protease separates CspB from CspA via CspBA interdomain processing^18,19,22^. Altering the length of this prodomain (due to loss in YabG or mutation) appears to alter *C. difficile*’s response to co-germinant signals^18^. Since the affinity of CspC for CspA is greater following CspBA interdomain processing (**Figure S1**), altering CspA’s N-terminal prodomain could impact CspA binding to CspC and thus its signaling function. Intriguingly, a similar subtilisin-like serine protease, PCSK9, uses an intrinsically disordered sequence in its N-terminal prodomain to allosterically regulate its affinity for lipoproteins vs. the low-density lipoprotein receptor (LDL-R)^40–44^. Future analyses of the role of CspA’s N-terminal prodomain on CspC:CspA heterodimer function could provide further insight into how this complex transduces germinant and co-germinant signals.

Finally, to address the third question, since pseudoenzymes frequently regulate their cognate enzymes by forming heterodimeric complexes with their active counterparts^34,35^, the CspC:CspA heterodimer could, in principle, activate the CspB protease via a “partner swap”, whereby one pseudoprotease would disassociate from the heterodimer and interact with CspB to regulate its activity. However, the extreme stability of the CspC:CspA heterodimer *in vitro*, both in the absence and presence of germinants and co-germinants, makes dissociation of the complex less likely (**Figures 4, 6, & 7**).

As an alternative to a “partner swap” model, pseudoenzymes can also serve as scaffolds that orchestrate the assembly and spatial organization of signaling complexes^34,35,45^. The germinant receptors in *B. subtilis* form germinosomes, where germination proteins spatially cluster^46^, so the CspC:CspA heterodimer may be similarly clustered near key germination proteins like CspB and SleC, potentially in partnership with GerG. By clustering germination components within a defined locus, the system could facilitate the efficient activation of CspB by the “active” conformation of the CspC:CspA heterodimer. Testing this possibility by localizing *C. difficile* germination proteins with the CspC:CspA heterodimer is the subject of further investigation.

The identification of the CspC:CspA heterodimer as a central signaling node during *C. difficile* germination may also inform studies of the germination mechanisms used by other clostridial organisms encoding Csp homologs. Notably, *C. perfringens* and *C. septicum* spores were recently shown to respond to bile acids^47,48^, and *C. septicum* CspC homologs have been genetically implicated in regulating bile acid-mediated germination^48^. Therefore, clostridial species may use a similar CspC:CspA heterodimer for inducing germination. Our finding that CspB forms a homodimer in *C. difficile* (**Figures 1 & S2**) just as in *C. perfringens*^19^ supports the notion that features of the germination machinery may be shared across a wide range of clostridial species. Future analyses of Csps and their interactions across other species will uncover the mechanisms underlying germination in the *Clostridia* and facilitate the development of strategies for inhibiting the germination of diverse clostridial pathogens.

## Materials and Methods

### Bacterial strains and growth conditions

*Escherichia coli* DH5α strains are listed in **Table S1** and were grown in Luria-Bertani (LB) broth at 37 °C with 225 rpm shaking. For protein production, *E. coli* BL21(DE3) strains were grown in autoinduction broth (Terrific broth [Thermo Fisher] supplemented with 0.5% glycerol, 0.05% glucose, and 0.1% α-lactose monohydrate). Media were supplemented with 20 μg/mL chloramphenicol, 100 μg/mL ampicillin, or 30 μg/mL kanamycin as needed.

*C. difficile* strains are listed in **Table S2** and were grown on brain heart infusion medium supplemented with 0.5% w/v yeast extract and 0.1% w/v L-cysteine (BHIS) with taurocholate (TA; 0.1% w/v; 1.9 mM), thiamphenicol (10-15 μg/mL), kanamycin (50 μg/mL), or cefoxitin (8 μg/mL) as needed. *C. difficile* defined medium (CDDM) was used for constructing complementation strains^49^. Cultures were grown swirling at 37 °C under anaerobic conditions using a gas mixture containing 85% N2, 5% CO2, and 10% H2.

### E. coli strain construction

All plasmids used in this study are compiled in **Table S1**, containing external links to plasmid maps with primer sequences used during cloning. *C. difficile* genes were codon-optimized (co) for *E. coli,* and plasmids were cloned via Gibson assembly and transformed into *E. coli* (DH5α). All plasmids were confirmed with whole plasmid sequencing by Plasmidsaurus using Oxford Nanopore Technology. Confirmed plasmids were transformed into *E. coli* HB101 for conjugation with *C. difficile* or *E. coli* BL21(DE3) for protein expression.

### Recombinant protein affinity purification

BL21(DE3) *E. coli* strains encoding Csps under the control of a P_lac_ promoter were grown in 20 mL LB with antibiotics as needed, back-diluted 1:1,000 into 1 L autoinduction broth with antibiotics as needed and grown for ∼60 hours at 20°C with 225 rpm shaking. Cultures were pelleted, resuspended in 50 mL low imidazole buffer (LIB; 500 mM NaCl, 50 mM Tris-HCl pH 7.5, 15 mM imidazole, 10% glycerol, 2 mM β-mercaptoethanol), and flash frozen in liquid nitrogen. Once thawed, cells were probe sonicated [Branson] in 3 x 45-second rounds at 40% amplitude with 5 minutes on ice between. Lysates were cleared by centrifugation at 1,000 rpm for 45 min at 4°C. Tagged proteins were affinity purified from cleared lysates using Ni-NTA agarose beads [Qiagen] with gentle rocking at 4 °C for 2 hours. The beads were washed three times with LIB before elution with either high imidazole buffer (HIB; 500 mM NaCl, 50 mM Tris-HCl pH 7.5, 200 mM imidazole, 10% glycerol, 2 mM β-mercaptoethanol) or by inducing cleavage of the CPD tag^27,28^ with 200 μM inositol hexakisphosphate (InsP_6_, Sigma) in LIB. Samples of whole cells before purification (induced), after lysis and removal of insoluble material (cleared lysate), and elution were analyzed by SDS-PAGE with Coomassie staining.

### Size exclusion chromatography

Affinity (or co-affinity) purified protein was buffer-exchanged into SEC buffer (200 mM NaCl, 10 mM Tris-HCl pH 7.5, 5% glycerol, 1 mM dithiothreitol) and concentrated using an Amicon Ultra-15 30 kDa cutoff centrifugal filter [Millipore Sigma]. Proteins were separated using a Superdex 200 Increase 10/300 GL column [GE] on an AKTA Pure high pressure liquid chromatography instrument [GE] with the following parameters: 500 μL loop volume, empty loop with 1 mL; 0.2 mL/min flow rate for 1 column volume. Apparent molecular weights were calculated using a BioRad gel filtration standard. Fractions were collected every 0.5 mL for SDS-PAGE analyses using Coomassie staining or Western blotting.

### CspA and CspC mixing experiments

CspA and CspC were purified as described above, mixed in a 1:1 ratio (w:w) in SEC buffer, and incubated for 2 hours at room temperature. The mixture was analyzed by SEC as described above. SEC fractions were analyzed by SEC followed by Western blotting. Western blots were imaged using LiCOR Odyssey instrument. Blots were quantified in LiCOR Image Studio software and each band was normalized to the sum of all bands in the respective channel.

### Protein purification for crystallography

Purified BL21(DE3) strains #3330 (**Table S1**) containing pET22b-CspC-CPD and pRSFDuet1-CspA(QS)_F944E_/_Y1092E_ plasmids, and strain #3127 (**Table S1**) containing pET22b-CspA-His_6_ were used for crystallography. For the case of the CspA homodimer, Ni-NTA agarose beads were eluted using high imidazole buffer. For the CspA:CspC heterodimer, instead of eluting the proteins using high imidazole, InsP_6_ was used to cleave the C-terminally CPD-His_6_ tag^27,28^ from CspC to purify the untagged version of the CspC-CspA heterodimer. Ni-NTA agarose beads were incubated with 200 µM InsP_6_ in LIB buffer with gentle shaking at 4 °C overnight. For both protein complexes, after the incubation with high imidazole or InsP_6_, Ni-NTA agarose beads were pelleted, and the supernatant collected. Pooled proteins were concentrated and injected into the Superdex 200 Increase 10/300 GL column. Fractions from the SEC peaks containing the pure CspA homodimer and CspA:CspC heterodimer, respectively, were collected and re-concentrated using Amicon Ultra-15 30 kDa cutoff centrifugal filter [Millipore Sigma].

### Crystallization and structure determination

#### CspA homodimer

Purified CspA homodimer, concentrated to 10 mg/mL, was mixed 1:1 with reservoir solution containing 0.2 M lithium sulfate, 0.1 M morpholinoethane sulfonic acid (MES) pH 6.0, and 20% w/v polyethylene glycol (PEG) 4000. Vapor diffusion in hanging drops yielded large stacks of rhomboidal plates after two weeks at room temperature. These crystal stacks were harvested and broken up for a microseed stock using glass beads. This microseed stock was diluted 1000,000-fold in reservoir solution and combined with soluble CspA at 4 mg/mL concentration. These conditions yielded individual rhomboidal plates between 50-100 μm across. Crystals were harvested and flash frozen using 20% glycerol as a cryoprotectant. X-ray diffraction data were collected at 100 °K using NE-CAT beam line ID-24-C at the Advanced Photon Source, Argonne National Laboratory, at a wavelength of 0.9786 Å. Data were preprocessed with the Xia2 pipeline using the Dials and Aimless modules. The initial structure of the CspA homodimer was phased by molecular replacement in PHASER using the CspC structure from *C. difficile* (PDBID: 6MW4)^25^ as an initial search model. The model was manually rebuilt in COOT^50^ and refined in PHENIX^51^ to a resolution of 3.2 Å. The atomic coordinates and structure factors were deposited to the RCSB Protein Data Bank under the accession number (PDBID): 9PR9.

#### CspC:CspA heterodimer

100 nL of purified CspC-CspA_EE_ heterodimer, concentrated to 10 mg/mL, was mixed 1:1 with reservoir solution containing 0.2 M magnesium chloride, 0.1 M Tris-HCl pH 8.5, and 20% w/v PEG 8000. Vapor diffusion in hanging drops yielded rhomboidal plates ∼100 μm in length after two weeks at room temperature. Crystals were harvested and flash frozen using 15% ethylene glycol as a cryoprotectant. X-ray diffraction data were collected at 100 °K using NE-CAT beam line ID-24-C at the Advanced Photon Source, Argonne National Laboratory, at a wavelength of 0.9786 Å. Data were preprocessed with the Xia2 pipeline^52^ using the Dials and Aimless modules. The initial structure of the CspC:CspA heterodimer was phased by molecular replacement in PHASER using the CspC structure from *C. difficile* (PDBID: 6MW4)^25^ as an initial search model. The model was manually rebuilt in COOT^50^ and refined in PHENIX^51^ to a resolution of 3.3 Å. The atomic coordinates and structure factors were deposited to the RCSB Protein Data Bank under the accession number (PDBID): 9PR8.

### C. difficile strain construction

Complementation strains were constructed as previously described^25,27,28^. Briefly, mutant *csp* genes were assembled in *pMTL-YN1C* plasmids containing the truncated *pyrE* gene^31,53^, cloned into DH5α cells, and sequence-confirmed using Plasmidsaurus. Sequence-confirmed plasmids were transformed into HB101 *E. coli* strains containing the necessary genes for conjugation. HB101 strains containing mutant *csp* complementation plasmids were combined with the associated *Δcsp C. difficile* strain to construct the *C. difficile* complementation strains. CDDM media was used to select for recombination of the construct into the *pyrE* locus, which restores the uracil prototrophy. Two independent clones from each conjugation were phenotypically characterized.

### Sporulation and spore purification

Sporulation and spore purification were performed as previously described^25,39^. Briefly, *C. difficile* strains were streaked from frozen glycerol stocks onto BHIS plates and grown overnight. Liquid BHIS cultures were grown until they reached an OD_600_ between 0.35 and 0.75. To induce sporulation, 100-120 μL of these cultures were spread onto 70:30 agar plates. Spores were collected from 70:30 plates a minimum of 60 hrs later and resuspended in ice-cold sterile water. To confirm proper levels of sporulation, a small sample was removed and examined via phase-contrast microscopy. Spore samples were washed with ice-cold water 5-8 times and incubated on ice overnight. The next day, the samples were treated with DNAse I [New England Biolabs] for 1 hr at 37 °C. After treatment, the spores were washed 1-2 times in ice-cold water and then purified on a 20%-50% Histodenz [Sigma Aldrich] gradient. After 2 more washes with water, spore purity was confirmed using phase-contrast microscopy. The samples were stored at 4 °C until use.

### OD_600_ kinetics assay

The OD_600_ kinetics assay protocol was adapted from previous works^11,25^. Spores were resuspended in BHIS media to an OD_600_ of 0.89 (∼7 x 10^5^ spore-forming units/μL), and 180 µL were added into wells of a 96-well flat-bottom tissue culture plate [Falcon] for each condition tested. The spores were exposed to different concentrations of TA (1%, 18.6 mM; 0.5%, 9.3 mM; 0.25%, 4.7 mM; 0.125%, 2.3 mM; 0.06%, 1.2 mM; 0.03%, 0.6 mM) or water (untreated) to a final volume of 200 µL per well. The OD_600_ was measured every 3 minutes using a Synergy H1 microplate reader [Biotek] at 37°C with constant shaking between readings. Readings were subtracted by background OD_600_ of spore-free media, and the values were normalized to the first measurements (time 0) and plotted as OD_600_ vs. time (**Figure S10**). Germination (arbitrary units) for each strain was calculated by first subtracting each of OD_600_ values from 1 to invert the curves (**Figure S10**). The area under the inverted curves was plotted for each strain as a bar graph (**Figure S10**). Area under the curve calculations were performed using Prism. All germination assays were initially performed using all 7 concentrations of TA to determine the concentration that best displays a given germinant sensitivity phenotype. This assay was performed for each strain at a minimum of three biological replicates, with three individually prepared spore isolates.

### Western blot analysis

Spore samples for immunoblotting were prepared as previously described^18,22,25^. Spores were diluted to an OD_600_ of 1.4 in 100 μL of sterile water (1.5 x 10^8^ spore-forming units), pelleted, and resuspended in 50 μL EBB buffer (8 M urea, 2 M thiourea, 4% (w/v) SDS, 2% (v/v) β-mercaptoethanol). The samples were boiled for 20 min with vigorous vortexing during incubation, pelleted, and resuspended. Final sample buffer with bromophenol blue was added to 1X to stain the sample. To maximally solubilize spore proteins, the samples were boiled again for 5-10 minutes, vortexed vigorously, and pelleted. The samples were resolved by 12% SDS-PAGE gels and transferred to Millipore Immobilon-FL PVDF membranes. The membranes were blocked with Odyssey Blocking Buffer with 0.1% (v/v) Tween 20 and probed with mouse monoclonal anti-CspC^22^ or anti-SleC^19^ and/or rabbit polyclonal anti-CspB^19^ or anti-CspA that was raised against SEC-purified CspA-His_6_ in rabbits (CoCalico Biologicals). The anti-CspC and anti-CspA antibodies were used at 1:2,000 dilutions, the anti-CspB antibody was used at 1:3,000, and the anti-SleC antibody was used at 1:5,000. The membranes were probed with primary antibodies for 4 hrs, washed 3x with PBS with 1.5x Tween 20, and probed with IRDye 680CW and 800CW infrared dye-conjugated secondary antibodies at 1:12,000. After incubation for 1hour in the dark, the membranes were washed twice with PBS with 1.5x Tween 20, and secondary antibody infrared fluorescence emissions were visualized using an Odyssey LiCOR CLx. The results shown are representative of three independently prepared spore isolates.

### Limited proteolysis

Limited proteolysis of purified *C. difficile* proteins was performed as previously described in *Rohlfing et al. 2019*. Purified *C. difficile* Csp proteins were diluted to 15 µM in 10 mM Tris pH 7.5 buffer. 24 µL of the protein solution was aliquoted into 0.2 mL PCR tubes. A 1 mg/mL chymotrypsin stock solution was diluted in 10 mM Tris pH 7.5 buffer to generate 10-fold dilutions. 1 µL of the appropriate chymotrypsin dilution was added to the protein solution to achieve the indicated concentration of chymotrypsin. 1 µL of 10 mM Tris pH 7.5 buffer was added to the protein samples as an untreated control. The protein and chymotrypsin solutions were incubated at 37°C for 1 hour. Chymotrypsin activity was then quenched with the addition of NuPAGE 4X LDS Sample Buffer [Invitrogen] and boiled for 3 min at 98°C. The samples were resolved on a 15% SDS-PAGE gel and visualized using Coomassie staining.

For limited proteolysis experiments with the addition of TA and glycine, the protein solutions and chymotrypsin were diluted as above. 23 µL of the protein solution was added to the 0.2 mL PCR tubes. A 10% (186 mM) TA stock solution and a 200 mM glycine stock solution were diluted to 12.5 mM in 10 mM Tris pH 7.5 buffer. 1 µL of the TA/glycine solution was added to the indicated protein samples for final concentrations of 0.5 mM TA and 0.5 mM glycine. 1 µL of the appropriate chymotrypsin dilution was added to the protein solution to achieve the indicated concentration of chymotrypsin. Incubation, protease quenching, resolution, and visualization were performed as described above.

### Data visualization

All graphs were prepared using GraphPad Prism. Gels and blots were imaged using a LiCOR Odyssey instrument. Protein structure visualization was performed using Pymol.

### Data and materials availability

Structure models and maps have been deposited to RCSB PDB with the following accession numbers: 9PR8 – CspC:CspA_EE_; and 9PR9 – CspA homodimer.

