## Supplementary tables and figures for "The CspC:CspA heterodimer transduces germinant and co-germinant signals during *Clostridioides difficile* spore germination"

**Supporting material:**

**Supplemental Table 1.** *E. coli* strains used in this study. Amino acid numbering based on YabG-cleaved CspA. "*cspBA* numbering" based on *cspBA* fusion gene. All *C. difficile* genes expressed in *E. coli* were designed with codon optimization for *E. coli* expression.

**Supplemental Table 2.** *C. difficile* strains used in this study. *cspBA* numbering based on *cspBA* fusion gene.

**Supplemental Table 3.** CspA homodimer and CspC:CspA heterodimer data collection and refinement. $CspA_{EE}$ = $CspA_{F944E\text{-}Y1092E}$ (*cspBA* fusion gene numbering), and $CspA_{F363E\text{-}Y511E}$ (YabG-cleaved CspA numbering).

**Supplemental Table 4.** PISA interface analyses of CspC-CspA heterodimer and CspA homodimer. PISA interface analysis identified hydrogen bond and salt bridge interactions at the hetero- and homodimeric interfaces. The heterodimer has a more extensive interface, with twenty hydrogen bonds and five salt bridges across the interface, whereas the CspA homodimer has fifteen hydrogen bonds and two salt bridges across the interface. Amino acid numbering based on YabG-cleaved CspA. "*cspBA* numbering" based on *cspBA* fusion gene.

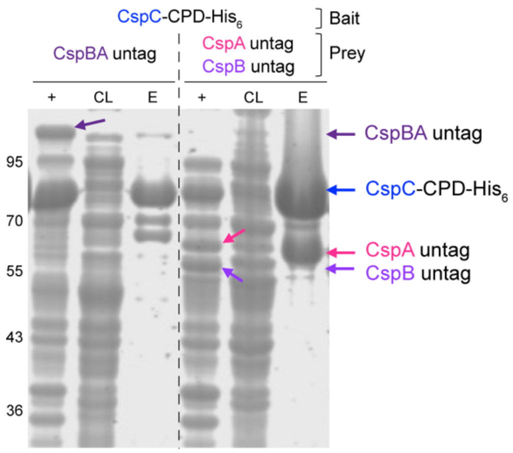


**Supplemental Figure 1. CspBA binds CspC with lower affinity than the CspA variant generated by interdomain processing.** Coomassie stain of co-affinity purifications using CspC-CPD-$His_6$ as the bait and untagged CspBA or CspB and CspA as the prey. +, induced fraction; CL, cleared lysate; E, elution. The data shown are representative of two independent replicates.

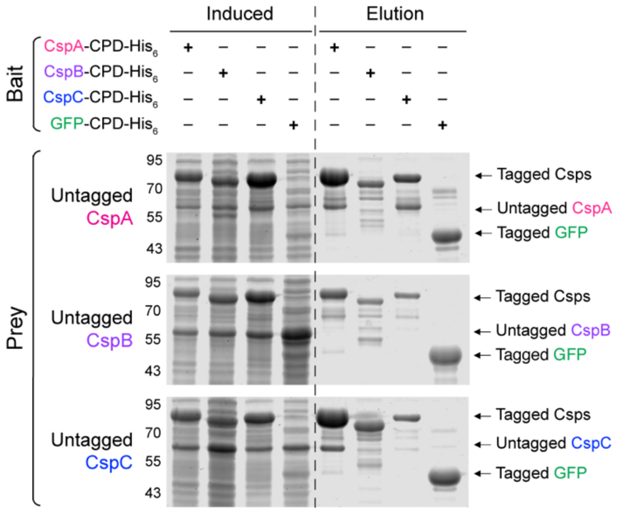


**Supplemental Figure 2. CspA and CspB form homodimers, but CspC does not.** Coomassie stain of co-affinity purification analyses using CPD-His$_6$-tagged CspA, CspB, or CspC as the bait alongside their respective untagged Csps as prey. GFP-CPD-His$_6$ is the control bait. The CspB prodomain (~9 kDa) is not shown. All data shown are representative of three replicates.

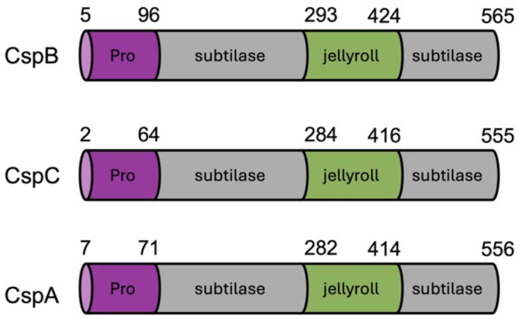


**Supplemental Figure 3. Csp domain boundaries.** Bounds of the prodomains, subtilase domains, and jellyroll domains of *C. perfringens* CspB (Adams *et al.*, 2013) (top), and *C. difficile* CspC and CspA (center and bottom, respectively). Bounds are representative of every structure in this work.

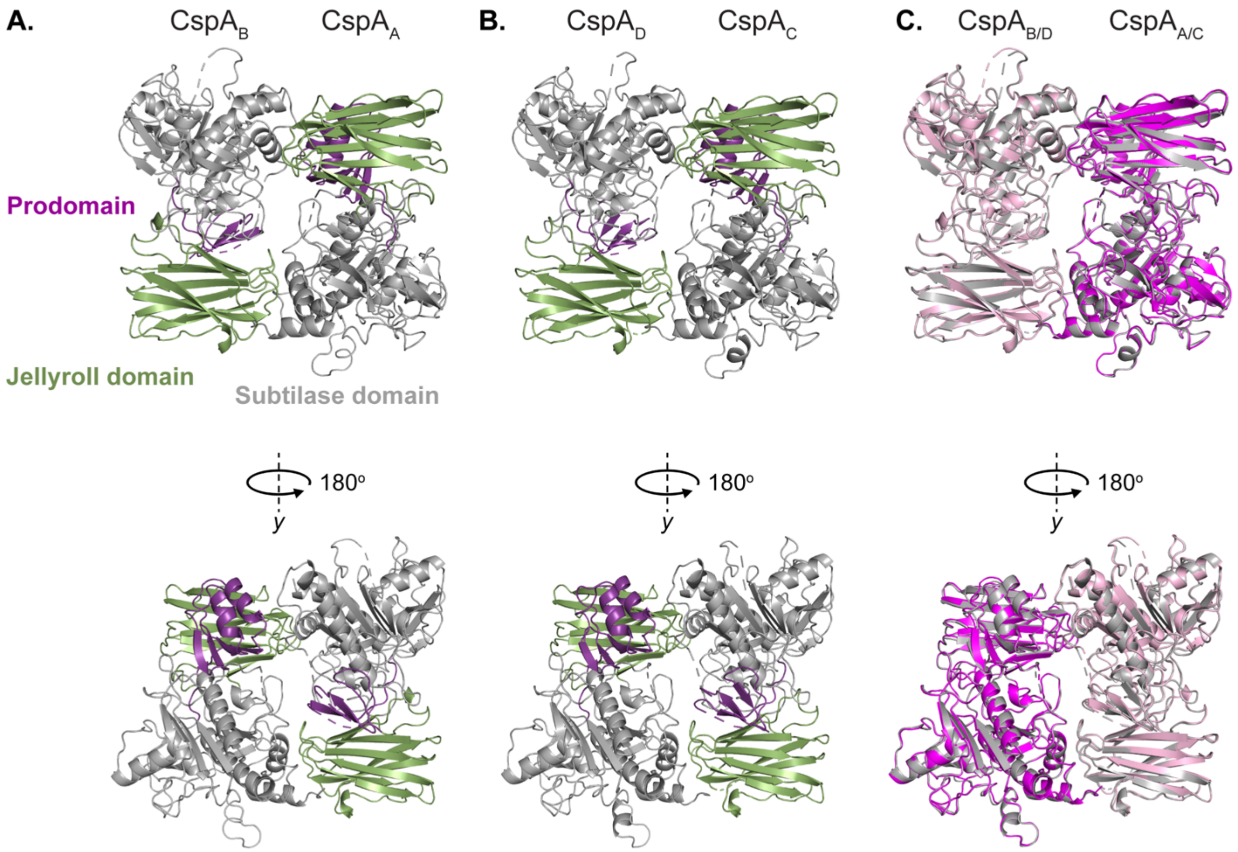


**Supplemental Figure 4. Asymmetric unit of CspA.** (A-B) The two CspA homodimers that crystallized within the asymmetric unit (PDB 9PR9). The subtilase domains are shown in grey, jellyroll domains in green, and prodomains in purple. The structure used for all other CspA homodimer figures in this manuscript is shown in (A). (C) Overlay of the two CspA homodimer structures (A & B). The A homodimer is shown in light pink ($CspA_B$) and magenta ($CspA_A$), the B homodimer is shown in grey.

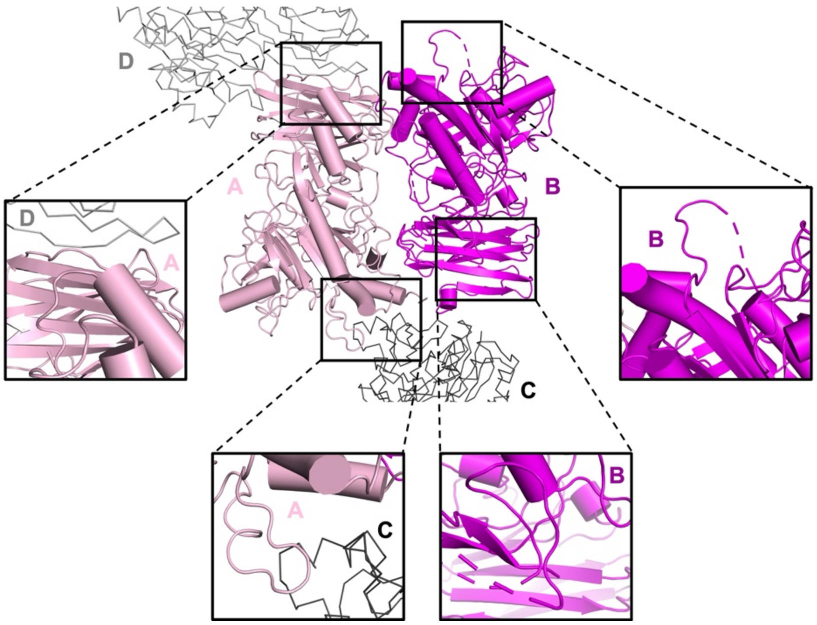


**Supplemental Figure 5. Crystal contacts with symmetry mates stabilize the CspA A and C protomer prodomain.** The prodomains of the A and C protomers in the CspA asymmetric unit are better resolved than their counterparts in the B and D protomers. As shown in the insets, the A protomer's prodomain makes contacts with C and D symmetry mates in the crystal lattice. The N terminus of the A prodomain is stabilized by contacts with the subtilase domain of a C protomer (bottom left inset), while the C terminus of the A prodomain is stabilized by contacts with the D protomer jellyroll domain (top right inset). Because of the screw axis and crystal packing, the B and D protomers lack these contacts with symmetry mates. Only protomers A and B of the asymmetric unit are shown for simplicity.

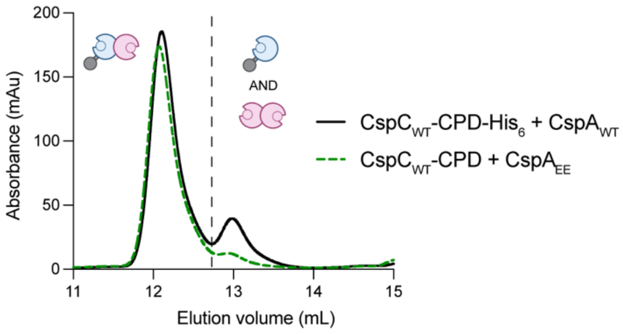


**Supplemental Figure 6. Size exclusion chromatography analyses of the protein variant used for CspC:CspA crystallization.** Size exclusion chromatography analysis of CspC-CPD-$His_6$-$CspA_{F944E/Y1092E}$ (CspC:$CspA_{EE}$) co-affinity purification. The dashed line indicates the separation between the two peaks. The data shown are representative of a minimum of three independent replicates.

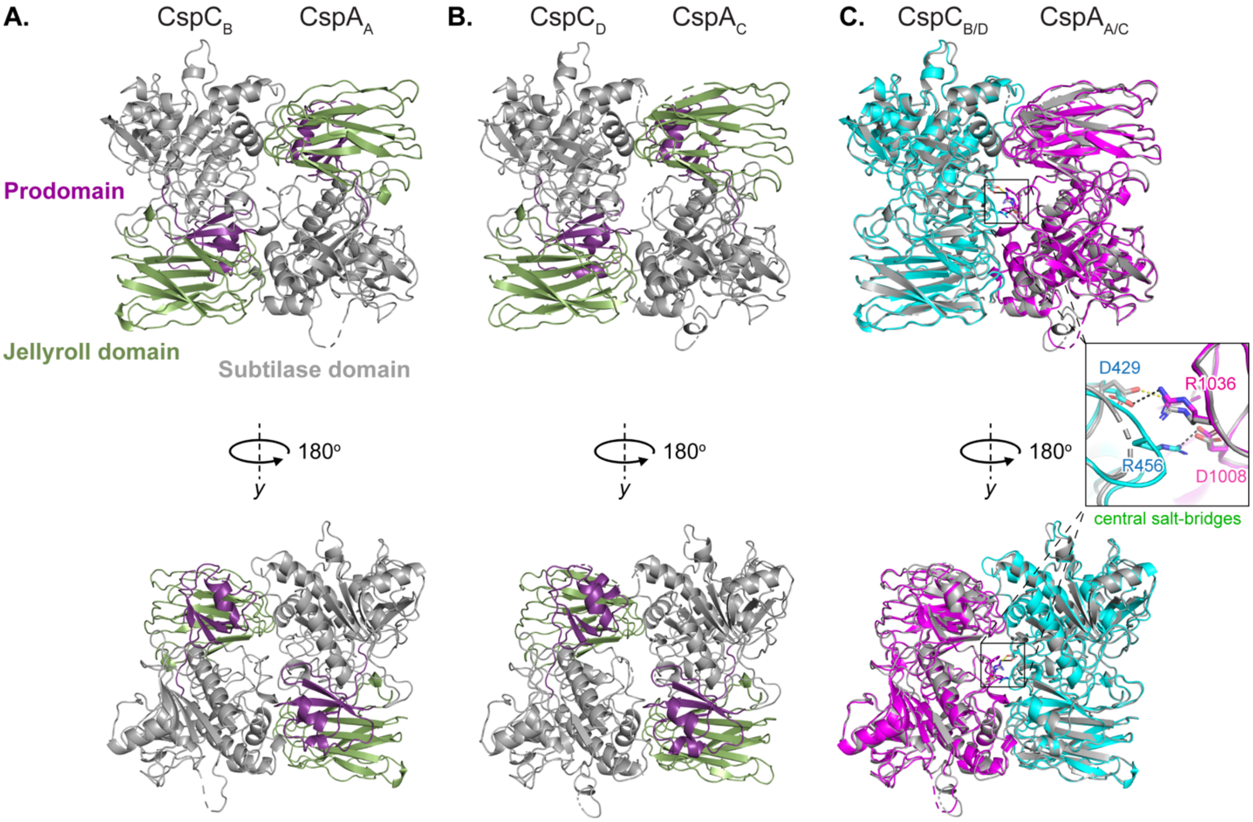


**Supplemental Figure 7. Asymmetric unit of CspC:CspA.** (A-B) The two CspC:CspA heterodimers that crystallized within the asymmetric unit (PDB 9PR8). CspC on the left, CspA on the right. The subtilase domains are shown in grey, jellyroll domains in green, and prodomains in purple. The structure used for all other CspC:CspA heterodimer figures in this manuscript is shown in (A). (C) Overlay of the two CspC:CspA heterodimer structures (A & B). The A heterodimer is shown in cyan ($CspC_B$) and magenta ($CspA_A$); the B heterodimer is shown in grey. (inset) Two salt bridges between CspC and CspA form within the $CspC_B$:$CspA_A$ heteterodimer. The $CspC_B$ D429:$CspA_A$ R1036 salt bridge interaction is a distance of 3.6 Å, and the $CspC_B$ R456:$CspA_A$ D1008 is 2.9 Å. A single salt-bridge forms between $CspC_D$ D429:$CspA_C$ R1036 (distance of 3.1 Å). The $CspC_D$ R456 residue is unstructured, and no interaction exists between $CspC_D$ R456 and $CspA_C$ D1008.

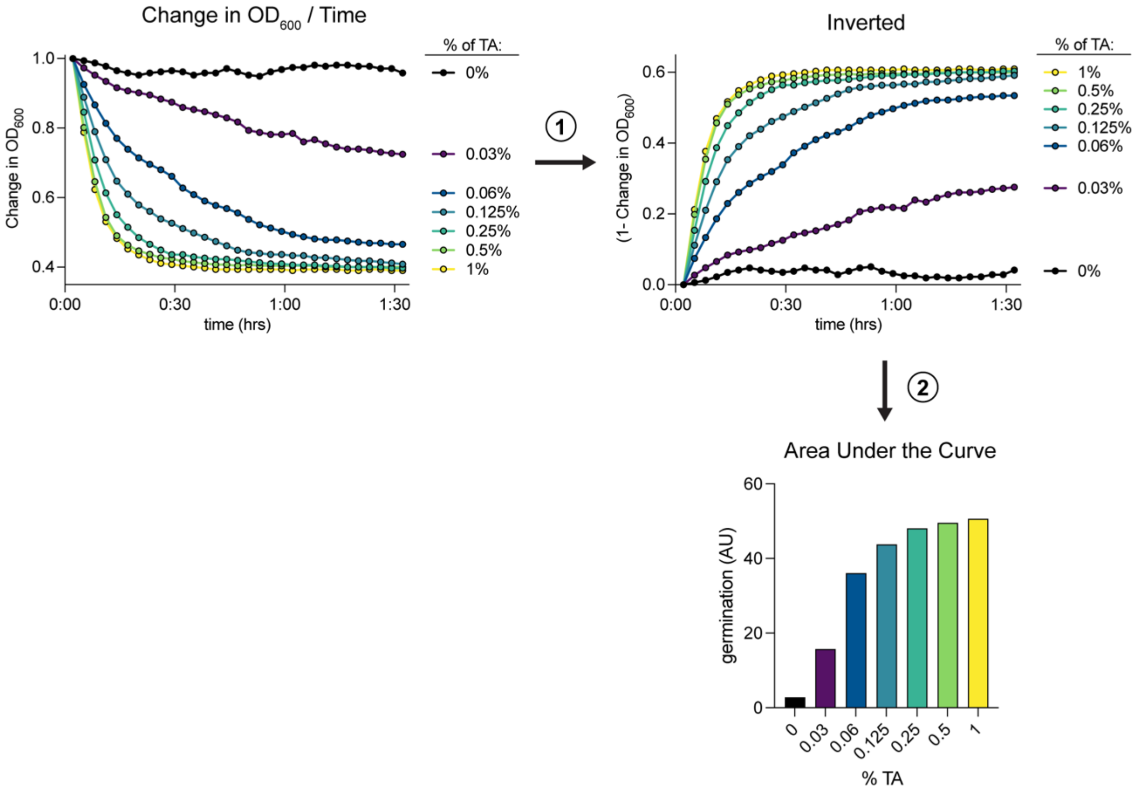


**Supplemental Figure 8. Schematic of $OD_{600}$ drop assay data presentation.** Spore germination was assessed using optical density ($OD_{600}$). Spores were exposed to germinant and their $OD_{600}$ was measured at 3-minute intervals for 1.5 hrs. After the $OD_{600}$ of blank wells (sterile medium) was subtracted, the values were normalized to the first measurements (time 0), and the values were plotted as $OD_{600}$ vs. time (left). (1) The resulting curves were inverted by subtracting each of the values from 1 (center). (2) The area under the inverted curves (center) was plotted for each strain as a bar graph (right). Area under the curve calculations were performed using Prism. All germination assays were performed at the presented concentrations of TA to determine the concentration that best displays a given germinant sensitivity phenotype.

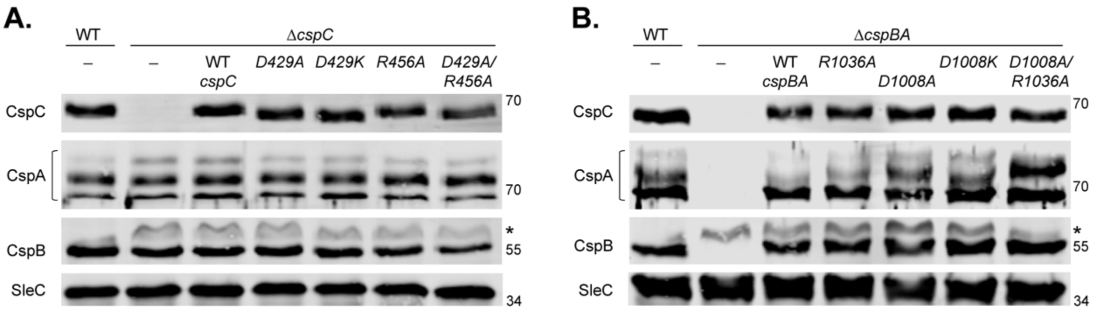


**Supplemental Figure 9. Mutation of two salt bridges at the CspC:CspA heterodimer interface does not impact Csp levels in mature spores.** (A-B) Western blot analyses of Csp levels in mutant spores. Multiple isoforms of CspA are observed. * indicates a non-specific band. SleC was used as a load control. The data shown are representative of a minimum of three independent replicates.

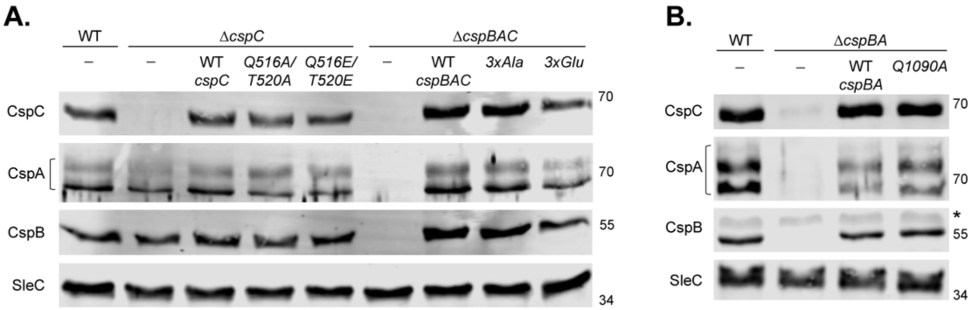


**Supplemental Figure 10. Mutations in peripheral H-bonds of the CspC:CspA heterodimer interface do not impact Csp levels in mature spores.** (A-B) Western blot analyses of Csp levels in mutant spores. Δ*cspBAC/3xAla* = triple-Ala substitution *cspBA*$_{R896A}$-*cspC*$_{Q516A/T520A}$, Δ*cspBAC/3xGlu* = triple-Glu substitution *cspBA*$_{R896E}$-*cspC*$_{Q516A/T520E}$. Multiple isoforms of CspA are observed. * indicates a non-specific band. SleC was used as a load control. The data shown are representative of a minimum of three independent replicates.

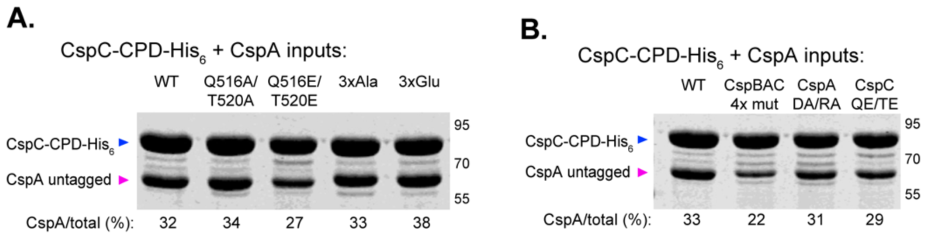


**Supplemental Figure 11. Inputs for CspC-CPD-His$_6$ + CspA SEC analyses.** (A-B) SDS-PAGE of CspC-CPD-His$_6$ co-affinity purification inputs for SEC purification, stained with Coomassie. Q516A/T520A represents CspC$_{Q516A/T520A}$-CPD-His$_6$ with untagged WT CspA; Q516A/T520E represents CspC$_{Q516E/T520E}$-CPD-His$_6$ with untagged WT CspA; 3xAla represents CspC$_{Q516A/T520A}$-CPD-His$_6$ with untagged CspA$_{R896A}$; and 3xGlu represents CspC$_{Q516E/T520E}$-CPD-His$_6$ with untagged CspA$_{R896E}$ (A). Inputs correspond to SEC traces shown in Figure 4E-G & K-L. (B) CspBAC 4x mut corresponds to CspC$_{Q516E/T520E}$-CPD-His$_6$ with untagged CspA$_{D1008A/R1036A}$; CspA DA/RA represents WT CspC-CPD-His$_6$ with untagged CspA$_{D1008A/R1036A}$; and CspC QE/TE represents CspC$_{Q516E/T520E}$-CPD-His$_6$ with untagged WT CspA. Inputs correspond to SEC traces shown in Figure 6D. CspA/total (%) = 100 x [CspA signal intensity / (CspA + CspC signal intensities)]. The data shown are representative of a minimum of two independent replicates.

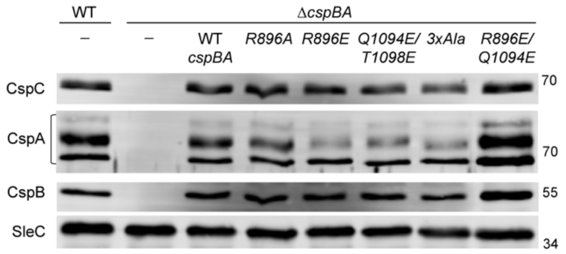


**Supplemental Figure 12. Mutations in the CspA homodimer twin H-bond networks do not impact Csp levels in mature spores.** Western blot analyses of Csp levels in mutant spores. *ΔcspBA/3xAla* = *ΔcspBA/cspBA*$_{R896A/Q1094A/T1098A.}$ Multiple isoforms of CspA are observed. * indicates a non-specific band. SleC was used as a load control. The data shown are representative of a minimum of three independent replicates.

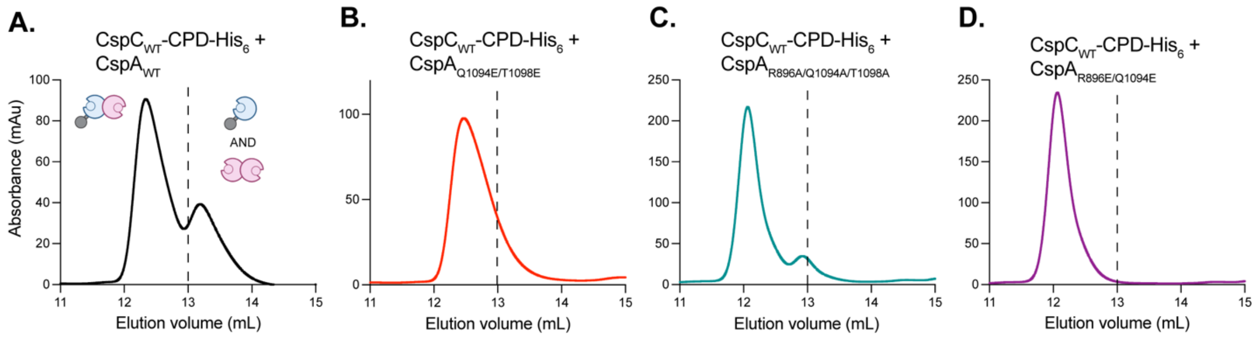


**Supplemental Figure 13. Effect of mutations in the CspA homodimer hydrogen bond network on CspC:CspA heterodimer formation.** Size exclusion chromatography analysis of CspC-CPD-His$_6$ and CspA variant co-affinity purifications. The dashed line indicates the separation between the two peaks. All data shown are representative of two independent replicates.

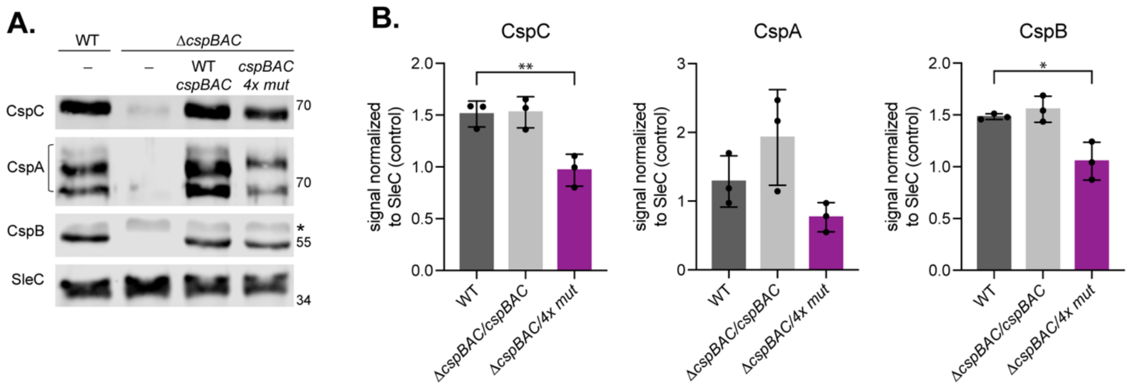


**Supplemental Figure 14. The CspC:CspA 4x mutant has decreased levels of Csps in purified spores.** (A) Western blot analyses of Csp levels in mutant spores. Δ*cspBAC/4X mut* = Δ*cspBAC/cspBA*$_{D1008A/R1036A}$-*cspC*$_{Q516E/T520E}$ Multiple isoforms of CspA are observed. Multiple isoforms of CspA are observed. * indicates a non-specific band. SleC was used as a load control. (B) Quantification of Western blot analyses. Protein signal intensities were normalized to the SleC loading control intensities. Statistical significance relative to WT was determined using a one-way ANOVA and Dunnett's multiple comparisons test. ** $p < 0.01$, * < 0.1. The data shown are representative of a minimum of three independent replicates.

**Supplementary Table 1**. *E.coli* strains used in this study

| Lab Strain # | Strain background | Genotype + plasmid carried | *cspBA* numbering | Source/Reference |
|---|---|---|---|---|
| 41 | DH5α | F– Φ80*lacZ* ΔM15 Δ(*lacZYA-argF*) U169 *recA1 endA1 hsdR17* (rK$^{-}$, mK$^{+}$) *phoA supE44* λ– *thi-1 gyrA96 relA1* | | D. cameron |
| 531 | HB101 | *F– mcrB mrr hsdS20(rB–mB–) recA13 leuB6 ara-13 proA2 lavYl galK2 xyl-6 mtl-1 rpsL20* carrying pRK24 | | C. Ellermeier |
| | BL21(DE3) | fhuA2 [lon] ompT gal (λ DE3) [dcm] ΔhsdS λ DE3 = λ sBamHIo ΔERI-B int::(lacI::PlacUV5::T7 gene1) i21 Δnin5 | | C2527 NEB |
| 3127 | BL21(DE3) | pET22b *cspA* -His$_6$ | | This study |
| 3179 | BL21(DE3) | pET22b cspB-His$_6$ | | This study |
| 981 | BL21(DE3) | pET22b *cspC* -His$_6$ | | *Rohlfing et al.* 2019. |
| 982 | BL21(DE3) | pET22b *cspC* -CPD-His$_6$ | | This study |
| 3125 | DH5α | pET22b *cspA* -CPD-His$_6$ | | This study |
| 3175 | DH5α | pET22b *cspB* -CPD-His$_6$ | | This study |
| 196 | DH5α | pET22b GFP-CPD-His$_6$ | | *Shen et al.* 2009. |
| 3028 | DH5α | pRSFDuet1 *cspA* | | This study |
| 3026 | DH5α | pRSFDuet1 *cspB* | | This study |
| 3177 | DH5α | pRSFDuet1 *cspC* | | This study |
| 3165 | BL21(DE3) | pRSFDuet1 *cspA* + pET22b *cspA* -CPD-His$_6$ | | This study |
| 3181 | BL21(DE3) | pRSFDuet1 *cspA* + pET22b *cspB* -CPD-His$_6$ | | This study |
| 3037 | BL21(DE3) | pRSFDuet1 *cspA* + pET22b *cspC* -CPD-His$_6$ | | This study |
| 3036 | BL21(DE3) | pRSFDuet1 *cspA* + pET22b GFP-CPD-His$_6$ | | This study |
| 3182 | BL21(DE3) | pRSFDuet1 *cspB* + pET22b *cspA* -CPD-His$_6$ | | This study |
| 3183 | BL21(DE3) | pRSFDuet1 *cspB* + pET22b *cspB* -CPD-His$_6$ | | This study |
| 3184 | BL21(DE3) | pRSFDuet1 *cspB* + pET22b *cspC* -CPD-His$_6$ | | This study |
| 3185 | BL21(DE3) | pRSFDuet1 *cspB* + pET22b GFP-CPD-His$_6$ | | This study |
| 3186 | BL21(DE3) | pRSFDuet1 *cspC* + pET22b *cspA* -CPD-His$_6$ | | This study |
| 3187 | BL21(DE3) | pRSFDuet1 *cspC* + pET22b *cspB* -CPD-His$_6$ | | This study |
| 3188 | BL21(DE3) | pRSFDuet1 *cspC* + pET22b *cspC* -CPD-His$_6$ | | This study |
| 3189 | BL21(DE3) | pRSFDuet1 *cspC* + pET22b GFP-CPD-His$_6$ | | This study |
| 3032 | DH5α | pRSFDuet1 *cspB* (SR)-*cspA* (QS) (separated at YabG cleavage site) | | This study |
| 3038 | BL21(DE3) | pRSFDuet1 *cspB* (SR)-*cspA* (QS) + pET22b GFP-CPD-His$_6$ | | This study |
| 1141 | DH5α | pET22b *cspC* -CPD-His$_6$ (SacI cleavage site) | | This study |
| 3039 | BL21(DE3) | pRSFDuet1 *cspB* (SR)-*cspA* (QS) + pET22b *cspC* -CPD-His$_6$ | | This study |
| 1375 | DH5α | pET28a *cspBA* | | This study |
| 3062 | BL21(DE3) | pET28a *cspBA* + pET22b *cspC* -CPD-His$_6$ | | This study |
| 3330 | BL21(DE3) | pRSFDuet1 cspA (*F363E/Y511E*) + pET22b *cspC* -CPD-His$_6$ | *cspBA (F944E/Y1092E)* | This study |
| 4498 | DH5α | pET22b *cspC* (*D429A/R455A*) -CPD-His$_6$ | | This study |
| 4501 | BL21(DE3) | pRSFDuet1 *cspA* (*F363E/Y511E*) + pET22b *cspC* (*D429A/R456A*) -CPD-His$_6$ | *cspBA (F944E/Y1092E)* | This study |
| 4497 | DH5α | pRSFDuet1 *cspA* (*D427A/R455A*) | *cspBA (D1008A/R1036A)* | This study |
| 4500 | BL21(DE3) | pRSFDuet1 *cspA* (*D427A/R455A*) + pET22b *cspC* -CPD-His$_6$ | *cspBA (D1008A/R1036A)* | This study |
| 3956 | DH5α | pET22b *cspC* (*Q516A/T520A*) -CPD-His$_6$ | | This study |
| 3959 | BL21(DE3) | pRSFDuet1 *cspA* + pET22b *cspC* (*Q516A/T520A*) -CPD-His$_6$ | | This study |
| 3441 | DH5α | pRSFDuet1 *cspA* (*R315A*) | *cspBA (R896A)* | This study |
| 4263 | DH5α | pET22b *cspC* (*Q516E/T520E*) -CPD-His$_6$ | | This study |
| 4267 | BL21(DE3) | pRSFDuet1 *cspA* + pET22b *cspC* (*Q516E/T520E*) -CPD-His$_6$ | | This study |
| 3443 | BL21(DE3) | pRSFDuet1 *cspA* (*R315A*) + pET22b cspC-CPD-His$_6$ | *cspBA (R896A)* | This study |
| 4355 | DH5α | pRSFDuet1 *cspA* (*R315E*) | *cspBA (R896E)* | This study |
| 4365 | BL21(DE3) | pRSFDuet1 *cspA* (*R315E*) + pET22b *cspC* -CPD-His$_6$ | *cspBA (R896E)* | This study |
| 3771 | DH5α | pRSFDuet1 *cspA* (*Q509A*) | *cspBA (Q1090A)* | This study |
| 3776 | BL21(DE3) | pRSFDuet1 *cspA* (*Q509A*) + pET22b *cspC* -CPD-His$_6$ | *cspBA (Q1090A)* | This study |
| 3962 | BL21(DE3) | pRSFDuet1 *cspA* (*R315A*) + pET22b *cspC* (*Q516A/T520A*) -CPD-His$_6$ | *cspBA (R896A)* | This study |
| 4429 | BL21(DE3) | pRSFDuet1 *cspA* (*R315E*) + pET22b *cspC* (*Q516E/T520E*) -CPD-His$_6$ | *cspBA (R896E)* | This study |
| 3420 | BL21(DE3) | pET22b *cspA* (*R315A*) -His$_6$ | *cspBA (R896A)* | This study |
| 4352 | BL21(DE3) | pET22b *cspA* (*R315E*) -His$_6$ | *cspBA (R896E)* | This study |
| 4266 | BL21(DE3) | pET22b *cspA* (*Q513A/T517A*) -His$_6$ | *cspBA (Q1094A/T1098A)* | This study |
| 3738 | BL21(DE3) | pET22b *cspA* (*R315E/Q513E*) -His$_6$ | *cspBA (R896E/Q1094E)* | This study |
| 3739 | BL21(DE3) | pET22b *cspA* (*R315A/Q513A/T517A*) -His$_6$ | *cspBA (R896A/Q1094A/T1098A)* | This study |
| 4497 | DH5α | pET22b *cspA* (*D427A-R455A*) -His$_6$ | *cspBA (D1008A/R1036A)* | This study |
| 4607 | BL21(DE3) | pRSFDuet1 cspA (*D427A/R455A*) + pET22b cspC (Q516E/T520E-CPD-His6 | *cspBA (D1008A/R1036A)* | This study |
| 4276 | DH5α | pET22b *cspA* (*Q513E/T517E*)-His$_6$ | *cspBA (Q1094E/T1098E)* | This study |
| 4279 | BL21(DE3) | pRSFDuet1 *cspA* (*Q513E/T517E*) + pET22b *cspC* -CPD-His$_6$ | *cspBA (Q1094E/T1098E)* | This study |
| 3775 | DH5α | pRSFDuet1 *cspA* (*R315A/Q513A/T517A*) | *cspBA (R896A/Q1094A/T1098A)* | This study |
| 3780 | BL21(DE3) | pRSFDuet1 *cspA* (*R315A/Q513A/T517A*) + pET22b *cspC* -CPD-His$_6$ | *cspBA (R896A/Q1094A/T1098A)* | This study |
| 3773 | DH5α | pRSFDuet1 *cspA* (*R315E/Q513E*) | *cspBA (R896E/Q1094E)* | This study |
| 3778 | BL21(DE3) | pRSFDuet1 *cspA* (*R315E/Q513E*) + pET22b *cspC* -CPD-His$_6$ | *cspBA (R896E/Q1094E)* | This study |
| 2319 | HB101 | pMTL-YN1C Δ*cspBA* -*cspC* (*D429A*) | | This study |
| 2323 | HB101 | pMTL-YN1C Δ*cspBA* -*cspC* (*R456A*) | | This study |
| 3415 | HB101 | pMTL-YN1C *cspBA* (*F944E/Y1092E*) -Δ*cspC* | | This study |
| 3464 | HB101 | pMTL-YN1C *cspBA* (*R896A*) -Δ*cspC* | | This study |
| 3631 | HB101 | pMTL-YN1C *cspBA* (*Q1094A*) -Δ*cspC* | | This study |
| 3756 | HB101 | pMTL-YN1C *cspBA* (*R896E/Q1094E*) -Δ*cspC* | | This study |
| 3763 | HB101 | pMTL-YN1C *cspBA* (*R896A/Q1094A/T1098A*) -Δ*cspC* | | This study |
| 3948 | HB101 | pMTL-YN1C Δ*cspBA* -*cspC* (*T520A*) | | This study |
| 3950 | HB101 | pMTL-YN1C Δ*cspBA* -*cspC* (Q516A/*T520A*) | | This study |
| 3977 | HB101 | pMTL-YN1C Δ*cspBA* -*cspC* (*Q516A*) | | This study |
| 4025 | HB101 | pMTL-YN1C *cspBA* (*R1036A*) -Δ*cspC* | | This study |
| 4140 | HB101 | pMTL-YN1C *cspBA* (*R896A*) *cspC* (*Q516A/T520A*) | | This study |
| 4246 | HB101 | pMTL-YN1C *cspBA* (*D1008K*) -Δ*cspC* | | This study |
| 4270 | HB101 | pMTL-YN1C Δ*cspBA* -*cspC* (*Q516E/T520E*) | | This study |
| 4327 | HB101 | pMTL-YN1C *cspBA* (*R896E*) -Δ*cspC* | | This study |
| 4329 | HB101 | pMTL-YN1C *cspBA* (*Q1094E/T1098E*) -Δ*cspC* | | This study |
| 4435 | HB101 | pMTL-YN1C *cspBA* (*R896E*) -*cspC* (*Q516E/T520E*) | | This study |
| 4437 | HB101 | pMTL-YN1C *cspBA* (*D1008A*) -Δ*cspC* | | This study |
| 4450 | HB101 | pMTL-YN1C *cspBA* (*D1008A/R1036A*) -Δ*cspC* | | This study |
| 4496 | HB101 | pMTL-YN1C Δ*cspBA* -*cspC* (*D429A/R456A*) | | This study |
| 4503 | HB101 | pMTL-YN1C *cspBA* (*D1008A/R1036A*) -cspC (*Q516E/T520E*) | | This study |

**Supplementary Table 2**. *C. difficile* strains used in this study

| Lab Strain # | Strain name | Relevant genotype | Source/Reference |
|---|---|---|---|
| 789 | 630Δ*erm*Δ*pyrE*Δ*cspBA* | *630*Δ*erm*Δ*pyrE* with *cspBA* deleted | *Kevorkian et al.* 2017. |
| 799 | 630Δ*erm*Δ*pyrE*Δ*cspC* | *630*Δ*erm*Δ*pyrE* with *cspC* deleted | *Kevorkian et al.* 2017. |
| 831 | 630Δ*erm*Δ*pyrE*Δ*cspC/cspC* | *630*Δ*erm*Δ*cspC* with *cspC* in the *pyrE* locus | *Kevorkian et al.* 2017. |
| 846 | 630Δ*erm*-P | *erm*-sensitive derivative of 630 with *pyrE* restored | *Kevorkian et al.* 2017. |
| 859 | 630Δ*erm*Δ*cspBA*-P | *630*Δ*erm*Δ*cspBA* with *pyrE* restored | *Kevorkian et al.* 2017. |
| 928 | 630Δ*erm*Δ*pyrE*Δ*cspBAC* | *630*Δ*erm*Δ*pyrE* with *cspBAC* deleted | *Kevorkian et al.* 2017. |
| 1150 | 630Δ*erm*Δ*cspBAC*-P | *630*Δ*erm*Δ*cspBAC* with *pyrE* restored | *Kevorkian et al.* 2017. |
| 1153 | 630Δ*erm*Δ*pyrE*Δ*cspBAC/cspBAC* | *630*Δ*erm*Δ*cspBAC* with *cspBAC* in the *pyrE* locus | *Donnelly et al.* 2017. |
| 1205 | 630Δ*erm*Δ*pyrE*Δ*cspBA/cspBA* | *630*Δ*erm*Δ*cspBA* with *cspBA* in the *pyrE* locus | *Kevorkian et al.* 2017. |
| 1239 | 630Δ*erm*Δ*cspC*-P | *630*Δ*erm*Δ*cspC* with *pyrE* restored | *Kevorkian et al.* 2017. |
| 1920 | 630Δ*erm*Δ*pyrE*Δ*cspC/cspC (D429K)* | *630*Δ*erm*Δ*cspC* with *cspC (D429K)* in the *pyrE* locus | *Rohlfing et al.* 2019. |
| 4168 | 630Δ*erm*Δ*pyrE*Δ*cspBA/cspBA (F944E/Y1092E)* | *630*Δ*erm*Δ*cspBA* with *cspBA (F944E/Y1092E)* in the *pyrE* locus | This study |
| 4241 | 630Δ*erm*Δ*pyrE*Δ*cspBA/cspBA (R896A)* | *630*Δ*erm*Δ*cspBA* with *cspBA (R896A)* in the *pyrE* locus | This study |
| 4377 | 630Δ*erm*Δ*pyrE*Δ*cspBA/cspBA (Q1090A)* | *630*Δ*erm*Δ*cspBA* with *cspBA (R896A)* in the *pyrE* locus | This study |
| 4499 | 630Δ*erm*Δ*pyrE*Δ*cspBA/cspBA (R896E/Q1094E)* | *630*Δ*erm*Δ*cspBA* with *cspBA (R896E/Q1094E)* in the *pyrE* locus | This study |
| 4617 | 630Δ*erm*Δ*pyrE*Δ*cspBA/cspBA (R896A/Q1094A/T1098A)* | *630*Δ*erm*Δ*cspBA* with *cspBA (R896A/Q1094A/T1098A)* in the *pyrE* locus | This study |
| 4847 | 630Δ*erm*Δ*pyrE*Δ*cspC/cspC (T520A)* | *630*Δ*erm*Δ*cspC* with *cspC (T520A)* in the *pyrE* locus | This study |
| 4850 | 630Δ*erm*Δ*pyrE*Δ*cspC/cspC (Q516A/T520A)* | *630*Δ*erm*Δ*cspC* with *cspC (Q516A/T520A)* in the *pyrE* locus | This study |
| 4877 | 630Δ*erm*Δ*pyrE*Δ*cspC/cspC (Q516A)* | *630*Δ*erm*Δ*cspC* with *cspC (Q516A)* in the *pyrE* locus | This study |
| 4880 | 630Δ*erm*Δ*pyrE*Δ*cspBA/cspBA (R1036A)* | *630*Δ*erm*Δ*cspBA* with *cspBA (R1036A)* in the *pyrE* locus | This study |
| 4975 | 630Δ*erm*Δ*pyrE*Δ*cspBAC/cspBA (R896A) cspC (Q516A/T520A)* | *630*Δ*erm*Δ*cspBAC* with *cspBA (R896A) cspC (Q516A/T520A)* in the *pyrE* locus | This study |
| 5077 | 630Δ*erm*Δ*pyrE*Δ*cspBA/cspBA (D1008K)* | *630*Δ*erm*Δ*cspBA* with *cspBA (D1008K)* in the *pyrE* locus | This study |
| 5091 | 630Δ*erm*Δ*pyrE*Δ*cspC/cspC (Q516E/T520E)* | *630*Δ*erm*Δ*cspC* with *cspC (Q516E/T520E)* in the *pyrE* locus | This study |
| 5147 | 630Δ*erm*Δ*pyrE*Δ*cspBA/cspBA (R896E)* | *630*Δ*erm*Δ*cspBA* with *cspBA (R896E)* in the *pyrE* locus | This study |
| 5150 | 630Δ*erm*Δ*pyrE*Δ*cspBA/cspBA (Q1094E/T1098E)* | *630*Δ*erm*Δ*cspBA* with *cspBA (Q1094E/T1098E)* in the *pyrE* locus | This study |
| 5276 | 630Δ*erm*Δ*pyrE*Δ*cspC/cspC (D429A)* | *630*Δ*erm*Δ*cspC* with *cspC (D429A)* in the *pyrE* locus | This study |
| 5278 | 630Δ*erm*Δ*pyrE*Δ*cspC/cspC (R456A)* | *630*Δ*erm*Δ*cspC* with *cspC (R456A)* in the *pyrE* locus | This study |
| 5287 | 630Δ*erm*Δ*pyrE*Δ*cspBAC/cspBA (R896E) cspC (Q516E/T520E)* | *630*Δ*erm*Δ*cspBAC* with *cspBA (R896E) cspC (Q516E/T520E)* in the *pyrE* locus | This study |
| 5328 | 630Δ*erm*Δ*pyrE*Δ*cspBA/cspBA (D1008A)* | *630*Δ*erm*Δ*cspBA* with *cspBA (D1008A)* in the *pyrE* locus | This study |
| 5330 | 630Δ*erm*Δ*pyrE*Δ*cspBA/cspBA (D1008A/R1036A)* | *630*Δ*erm*Δ*cspBA* with *cspBA (D1008A/R1036A)* in the *pyrE* locus | This study |
| 5451 | 630Δ*erm*Δ*pyrE*Δ*cspC/cspC (D429A/R456A)* | *630*Δ*erm*Δ*cspC* with *cspC (D429A/R456A)* in the *pyrE* locus | This study |
| 5455 | 630Δ*erm*Δ*pyrE*Δ*cspBAC/cspBA (D1008A/R1036A) cspC (Q516E/T520E)* | *630*Δ*erm*Δ*cspBAC* with *cspBA (D1008A/R1036A) cspC (Q516E/T520E)* in the *pyrE* locus | This study |

| Supplemental Table S3. Crystallographic data collection and refinement statistics* | | |
|---|---|---|
| **Construct (PDB ID)** | **CspA homodimer (9PR9)** | **CspC-CspA$_{EE}$ (9PR8)** |
| **Data Collection** | | |
| Resolution range (Å) | 100.6 - 3.22 (3.3 - 3.22) | 99.92 - 3.35 (3.45 - 3.35) |
| Space group | P $2_1$ $2_1$ $2_1$ | P $2_1$ $2_1$ $2_1$ |
| Unit cell (Å)<br>α (°) | 125.98 131.93 155.327<br>90 | 71.143 105.505 311.325<br>90 |
| Multiplicity | 7.9 (8.2) | 12.3 (11.9) |
| Completeness (%) | 98.23 (86.37) | 99.67 (97.65) |
| Mean I/sigma(I) | 3.1 (0.6) | 3.6 (0.8) |
| R-merge | 0.64 (3.83) | 0.87 (4.21) |
| R-meas | 0.68 (4.09) | 0.91 (4.39) |
| R-pim | 0.24 (1.41) | 0.26 (1.25) |
| $CC_{1/2}$ | 0.97 (0.33) | 0.97 (0.37) |
| **Refinement** | | |
| Reflections used in refinement | 41835 (2572) | 34565 (2745) |
| Reflections used for R-free | 1991 (122) | 1733 (145) |
| R-work | 0.2724 (0.3816) | 0.2457 (0.3531) |
| R-free | 0.3190 (0.4054) | 0.2777 (0.3761) |
| Number of non-hydrogen atoms | 15717 | 15999 |
| macromolecules | 15717 | 15997 |
| ligands | 0 | 0 |
| solvent | 0 | 2 |
| Protein residues | 2076 | 2165 |
| $RMS_{bonds}$ (Å) | 0.005 | 0.011 |
| $RMS_{angles}$ (°) | 1.08 | 1.36 |
| Average B-factor | 67.75 | 81.22 |
| macromolecules | 67.75 | 81.23 |
| ligands | | |
| solvent | | 68.84 |

***Values in parentheses are for highest-resolution shell.**

**Supplementary Table S4. PISA analysis of interface interactions.**

| CspC:CspA Heterodimer | | | | |
|---|---|---|---|---|
| | Chain:Residue # [Atom] | Distance (Å) | Chain:Residue # [Atom] | CspBA numbering |
| Hydrogen Bonds | | | | |
| 1 | C:ARG 510[ NE ] | 3.80 | A:GLN 55[ O ] | 636 |
| 2 | C:ARG 510[ NH2] | 3.45 | A:LEU 56[ O ] | 637 |
| 3 | C:ARG 510[ NH2] | 3.66 | A:ILE 59[ O ] | 640 |
| 4 | C:ASN 262[ ND2] | 2.85 | A:SER 256[ OG ] | 837 |
| 5 | C:TYR 521[ OH ] | 3.49 | A:ASN 338[ OD1] | 919 |
| 6 | C:GLN 516[ NE2] | 3.33 | A:GLU 363[ O ] | 944 |
| 7 | C:THR 520[ OG1] | 2.90 | A:GLY 365[ O ] | 946 |
| 8 | C:LYS 528[ NZ ] | 3.54 | A:TYR 389[ OH ] | 970 |
| 9 | C:LYS 319[ NZ ] | 3.17 | A:VAL 549[ O ] | 1130 |
| 10 | C:TYR 337[ OH ] | 2.93 | A:LEU 550[ O ] | 1131 |
| 11 | C:ILE 50[ O ] | 3.13 | A:ARG 265[ NH2] | 846 |
| 12 | C:VAL 52[ O ] | 3.83 | A:ARG 265[ NH2] | 846 |
| 13 | C:ALA 53[ O ] | 3.86 | A:ARG 265[ NE ] | 846 |
| 14 | C:TYR 361[ O ] | 2.76 | A:GLN 509[ NE2] | 1090 |
| 15 | C:GLY 367[ O ] | 2.69 | A:THR 517[ OG1] | 1098 |
| 16 | C:ASP 429[ OD2] | 3.39 | A:ARG 455[ NH1] | 1036 |
| 17 | C:ASP 429[ OD2] | 3.56 | A:ARG 455[ NH2] | 1036 |
| 18 | C:LEU 511[ O ] | 3.39 | A:ASN 338[ ND2] | 919 |
| 19 | C:GLN 516[ OE1] | 2.65 | A:ARG 315[ NH2] | 896 |
| 20 | C:THR 520[ OG1] | 3.06 | A:ARG 315[ NH1] | 896 |
| Salt Bridges | | | | |
| 1 | C:ARG 456[ NH1] | 3.95 | A:ASP 427[ OD1] | 1008 |
| 2 | C:ARG 456[ NH2] | 3.84 | A:ASP 427[ OD1] | 1008 |
| 3 | C:ARG 456[ NH2] | 2.87 | A:ASP 427[ OD2] | 1008 |
| 4 | C:ASP 429[ OD2] | 3.39 | A:ARG 455[ NH1] | 1036 |
| 5 | C:ASP 429[ OD2] | 3.56 | A:ARG 455[ NH2] | 1036 |

| CspA Homodimer | | | | | |
|---|---|---|---|---|---|
| | CspBA numbering | Chain:Residue # [Atom] | Distance (Å) | Chain:Residue # [Atom] | CspBA numbering |
| Hydrogen Bonds | | | | | |
| 1 | 846 | B:ARG 265[ NH2] | 3.11 | A:GLN 55[ OE1] | 636 |
| 2 | 1098 | B:THR 517[ OG1] | 2.77 | A:GLY 365[ O ] | 946 |
| 3 | 896 | B:ARG 315[ NH1] | 3.81 | A:GLN 513[ O ] | 1094 |
| 4 | 896 | B:ARG 315[ NH2] | 2.57 | A:GLN 513[ OE1] | 1094 |
| 5 | 896 | B:ARG 315[ NH1] | 3.13 | A:THR 517[ OG1] | 1098 |
| 6 | 970 | B:TYR 389[ OH ] | 3.88 | A:VAL 549[ O ] | 1130 |
| 7 | 899 | B:LYS 318[ NZ ] | 3.27 | A:GLU 553[ OE2] | 1134 |
| 8 | 915 | B:VAL 334[ O ] | 2.88 | A:HIS 554[ NE2] | 1135 |
| 9 | 919 | B:ASN 338[ O ] | 3.81 | A:HIS 555[ N ] | 1136 |
| 10 | 920 | B:TYR 339[ OH ] | 3.85 | A:HIS 556[ ND1] | 1137 |
| 11 | 946 | B:GLY 365[ O ] | 3.15 | A:THR 517[ OG1] | 1098 |
| 12 | 1094 | B:GLN 513[ O ] | 3.69 | A:ARG 315[ NH1] | 896 |
| 13 | 1094 | B:GLN 513[ OE1] | 2.30 | A:ARG 315[ NH2] | 896 |
| 14 | 1098 | B:THR 517[ OG1] | 2.95 | A:ARG 315[ NH1] | 896 |
| 15 | 1131 | B:LEU 550[ O ] | 3.36 | A:ASN 338[ ND2] | 919 |
| Salt Bridges | | | | | |
| 1 | 899 | B:LYS 318[ NZ ] | 3.80 | A:GLU 553[ OE1] | 1134 |
| 2 | 899 | B:LYS 318[ NZ ] | 3.27 | A:GLU 553[ OE2] | 1134 |
